# Cellular adaptation of mitochondrial metabolism and biogenesis in response to partial ATP synthase uncoupling

**DOI:** 10.64898/2025.12.18.695256

**Authors:** Elodie Sardin, François Godard, Jean-Paul Lasserre, Bénédicte Salin, Laurianne Daniel, Jenny Renaut, Jean-Paul di Rago, Stéphane Duvezin-Caubet, Emmanuel Tetaud

## Abstract

Cellular energy metabolism relies on tight regulation of mitochondrial biogenesis and turnover to maintain bioenergetic homeostasis. Understanding the signaling pathways that modulate mitochondrial content in response to changing energetic demands is therefore essential. This work examines how partial mitochondrial uncoupling reshapes mitochondrial content and metabolism in yeast and how such adaptations mirror features of pathological or age-associated states. Using a *Saccharomyces cerevisiae* ATP synthase mutant lacking the epsilon subunit (Δε) and carrying a compensatory mutation that allows survival under uncoupling (Δε+c-L57F), mitochondrial abundance, organization, and function were compared with those of wild-type and rescue strains (mutant re-expressing the epsilon subunit of the ATP synthase). Analyses included respiration assays, mitochondrial DNA quantification, β-galactosidase reporters, fluorescence microscopy of the mitochondrial network, and 2D-DiGE proteomics. Uncoupling in the mutant led to a strong increase in mitochondrial mass per cell, higher respiratory capacity, elevated cytochrome levels, and increased mtDNA copy number, together with augmented transcriptional activity consistent with activation of HAP-dependent mitochondrial biogenesis. Despite a fragmented network, mitochondria retained typical ultrastructural features, while proteomic profiling indicated a shift toward ATP production via substrate-level phosphorylation. Overall, partial uncoupling promotes reversible mitochondrial biogenesis and metabolic remodeling that depend on a preserved proton-motive force, highlighting mitochondrial signaling as a key regulator of cellular energy homeostasis and offer new perspectives for studying metabolic disorders and mitochondrial pathologies.

**GRAPHICAL ABSTRACTS:** 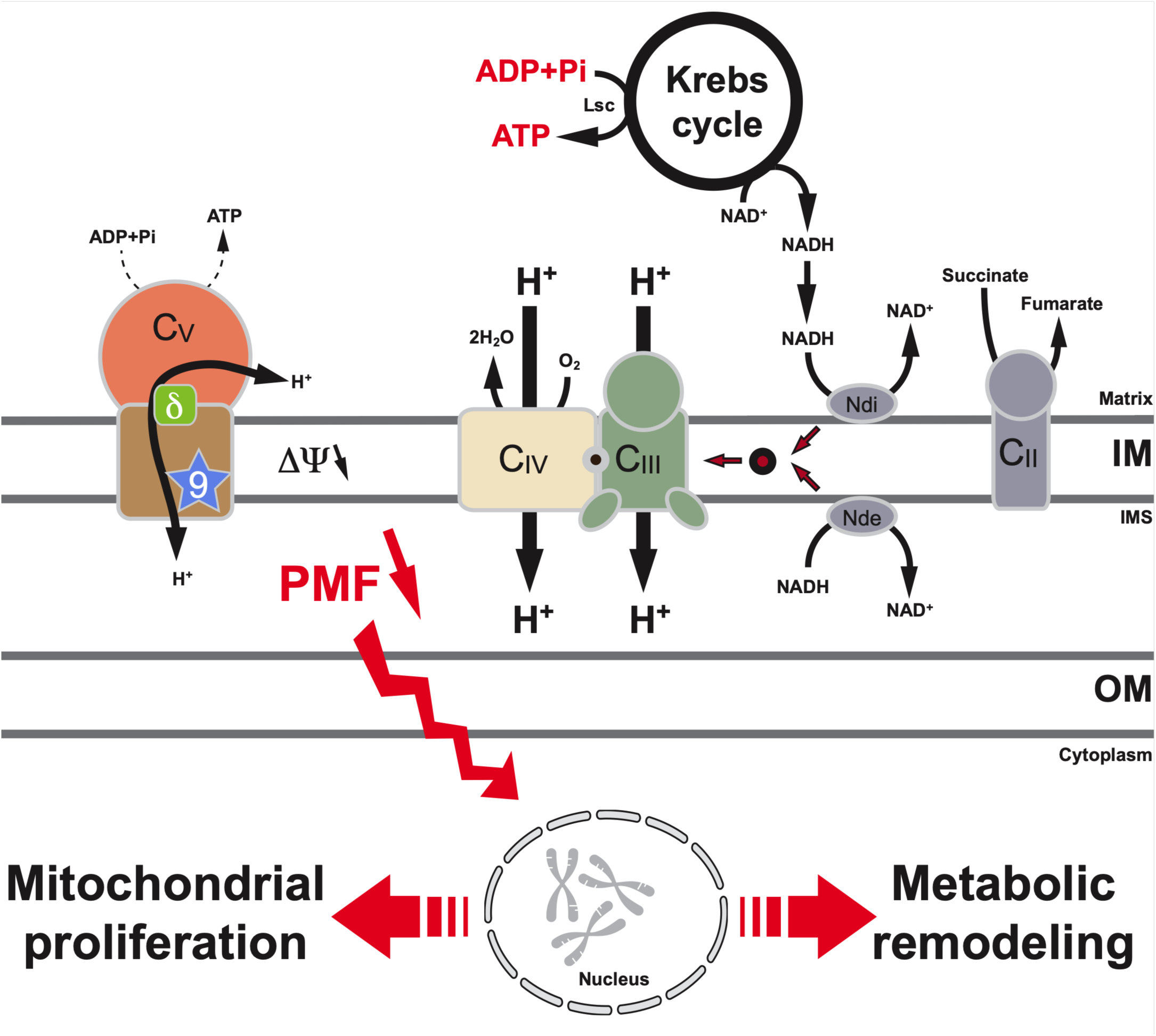

**Highlights:**

- Mitochondrial uncoupling triggers a marked increase in mitochondrial biogenesis in yeast.
- Uncoupling leads to fragmentation of the mitochondrial network, while preserving mitochondrial ultrastructure.
- Uncoupling induces metabolic remodeling that favors ATP production through substrate-level phosphorylation.
- These cellular adaptations are reversible upon re-establishment of mitochondrial coupling.
- The proton motive force serves as a key signal sensed by the cell.

## 1. INTRODUCTION

Various diseases such as diabetes, cardiovascular diseases, neurodegenerative disorders, muscular dystrophies, and cancers exhibit significant deregulation of cellular metabolism [1–4]. Understanding the mechanisms that govern cellular bioenergetic homeostasis, particularly mitochondrial homeostasis, is crucial for energy metabolism. Research in mammals and yeast has shown multiple ways to adjust mitochondrial metabolism to meet cellular energy demands, either by modulating oxidative phosphorylation kinetics/efficiency or altering mitochondrial enzymatic content [5]. Two key opposing processes, mitochondrial biogenesis and autophagy/mitophagy, finely tune mitochondrial homeostasis, with deregulation of either leading to functional deterioration of biological systems, potentially resulting in cell death [6; 7].

Mitochondrial biogenesis is a complex process jointly regulated by the mitochondrial and nuclear genomes [8; 9]. In mammalian cells, central to this regulation are the nuclear respiratory factors (NRF), which control the transcription of genes encoding subunits of the mitochondrial respiratory chain complexes [10–12]. The peroxisome proliferator-activated receptor γ coactivators 1 (PGC-1) family also plays a crucial role, with PGC-1α primarily driving mitochondrial biogenesis, metabolic responses, and enhanced mitochondrial respiration, while PGC-1β maintains basic mitochondrial function [2; 13–16]. In yeast, *Saccharomyces cerevisiae*, mitochondrial biogenesis is regulated by the Hap (heme activating proteins) complex, which includes Hap2, 3, 4, and 5, with Hap4 acting as the coactivator and functional homologue of mammalian PGC-1α [17–21]. This complex regulates the expression of genes involved in the Krebs cycle and the oxidative phosphorylation system (OXPHOS) [22]. Contrasting biogenesis, autophagy and mitophagy (selective autophagy) degrade mitochondria, regulating mitochondrial mass and ensuring mitochondrial quality control. These mechanisms facilitate the selection of damaged mitochondrial components for lysosomal degradation [7; 23–29].

While we are beginning to understand the mechanisms and actors involved in mitochondrial biogenesis regulation, the signaling pathways that control mitochondrial biogenesis are less well-characterized. Studies in mammalian cells and yeast demonstrate that cyclic adenosine monophosphate (cAMP) plays a role in inducing mitochondrial biogenesis [30]. Additionally, research in yeast highlights reactive oxygen species (ROS) as down-regulators of mitochondrial biogenesis, whereas in mammalian cells, mitochondrial ROS can either increase or decrease mitochondrial biogenesis [7].

Significant insights into mitochondrial functions and pathogenic mechanisms of mitochondrial diseases have been gained using the yeast model *Saccharomyces cerevisiae* [31]. This unicellular organism shares a high conservation of mitochondrial functions, allows for manipulation of both nuclear and mitochondrial genomes, and facilitates extensive genetic and chemical screenings. In this study, we explore mitochondrial biogenesis regulation in response to cellular energy demands using a yeast ATP synthase mutant that survives mitochondrial uncoupling [32]. Our findings demonstrate significant stimulation of mitochondrial biogenesis and precise reprogramming of energy metabolism, highlighting the pivotal role of the proton-motive force in mitochondrial energy metabolism adaptations. These results provide a foundation for further studies on energy metabolism adaptations during development, cellular differentiation, environmental changes, or metabolic disorders observed in various pathologies.

## 2. RESULTS

### 2.1. Mitochondrial uncoupling leads to a massive mitochondrial biogenesis

#### 2.1.1. Influence of the mitochondrial uncoupling on the amount of mitochondria

To investigate how mitochondrial biogenesis is regulated in response to cellular energy demands, we used a yeast mutant deficient in mitochondrial ATP synthase activity and capable of surviving mitochondrial uncoupling. This mutant, previously described in our laboratory [32], demonstrates that deletion of the ε-subunit of mitochondrial ATP synthase leads to complete mitochondrial uncoupling that can be partially compensated by a point mutation (L57F) in the c-subunit of the enzyme. In this study, three yeast strains will be examined in parallel: (i) the strain YE1, which expresses the ε-subunit under a doxycycline-regulatable promoter and behaves as a wild-type strain in the absence of doxycycline, but shows total mitochondrial uncoupling in its presence; (ii) the mutant S1, which lacks the ε-subunit but has the c-subunit mutation (c-L57F) leading to partial mitochondrial uncoupling and enables cells survival; and (iii) the strain S1+ε, which re-expresses the ε-subunit in the S1 mutant background, serving as a rescue strain [32]. As expected, the YE1, S1 (Δε+c-L57F), and S1+ε strains exhibited robust growth on fermentable carbon sources, such as glucose (Figure 1A). However, the S1 strain growth was significantly impaired on respiratory carbon sources like glycerol, while the S1+ε strain showed growth comparable to the wild-type YE1 on these carbon sources (Figure 1B) demonstrating that the c-L57F mutation does not affect growth under conditions where mitochondria remain coupled.

**Figure 1 -.**
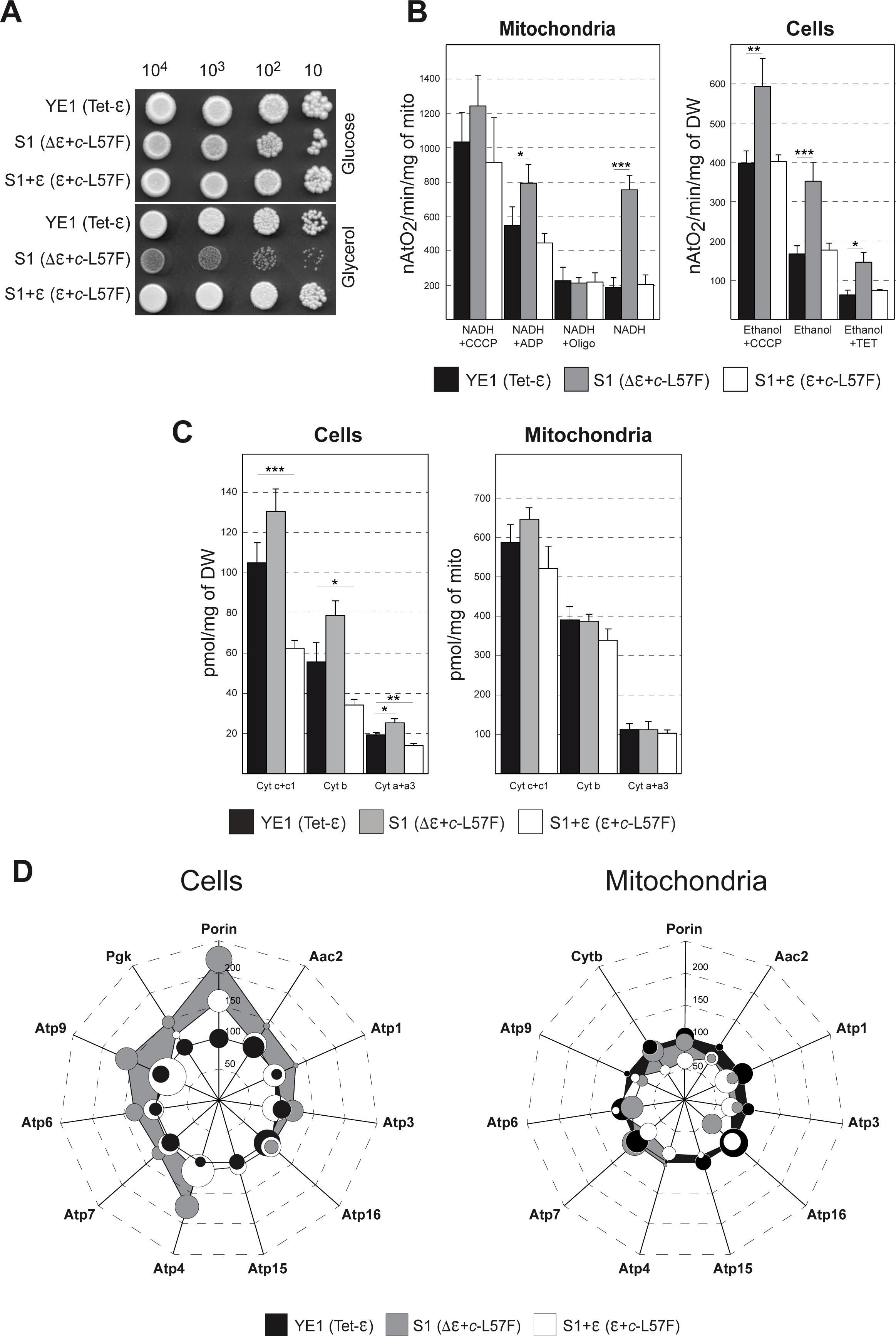
Cells and mitochondria properties of the YE1, S1 and S1+ε strains. **(A)** Drop tests of YE1 (Tet-ε), S1 (Δε+*c*-L57F) and S1+ε (ε+*c*-L57F) strains on glucose or glycerol and grown for 3-5 days at 28 °C. **(B)** Oxygen consumption of YE1 (black bars), S1 (gray bars) and S1+ε (white bars) isolated mitochondria or cells. The uncoupled respiratory rate was assessed in the presence of 5 µM carbonyl cyanide p-chlorophenylhydrazone (CCCP), the phosphorylation rate in the presence of NADH+ADP (Mitochondria) or ethanol (Cells) and the respiratory rate in the absence of phosphorylation in the presence of 20 µg/mg oligomycin (Oligo, mitochondria) or 50 µM triethyltin (TET, cells; [102; 103]). DW, dry weight. **(C)** Cytochrome content of YE1 (black bars), S1 (gray bars) and S1+ε (white bars) cells or isolated mitochondria. DW, dry weight. **(D)** Quantifications from immunoblot of several ATP synthase subunits, ADP/ATP carrier (Aac2), phosphoglycerate kinase (Pgk) in total cell extracts and cytochrome b (Cytb) in isolated mitochondria, normalized against porin abundance of the YE1 strain grown in 2% glycerol/ethanol medium at 28°C. Data are represented as a « Radar chart » where YE1 strain (black) was setup a 100% and compared to S1 (gray) and S1+ε (white). Mean ± SEM are represented by small circles on the radar chart. Data are represented as mean ± SEM of at least three independent cell cultures and mitochondria preparations. See also Supplemental Figure 1A/B/C/D.

Biochemical analysis of purified mitochondria from glycerol/ethanol-grown YE1, S1 (Δε+c-L57F) and S1+ε strains was previously reported [32] but to aid in understanding the following results, these analyses are presented again (Figure 1B, Mitochondria), but this time, compared with similar analyses on whole cells (Figure 1B, Cells). Mitochondrial oxygen consumption was measured using NADH as an electron donor under various conditions: with the membrane potential uncoupler CCCP (NADH+CCCP, representing maximum respiration), under phosphorylating conditions (NADH+ADP, state 3), with oligomycin (NADH+Oligo, a specific inhibitor of ATP synthase’s proton-translocating domain, state 4), and under basal respiration (state 4 without ADP). Compared to YE1, S1 showed a slight increase in maximum respiratory activity (NADH+CCCP). As previously shown [32], S1 exhibited a partial mitochondrial uncoupling, blocked by oligomycin (Figure 1B, Mitochondria, NADH vs NADH+Oligo). CCCP efficiently stimulated respiration in all strains, indicating normal passive permeability to protons in the mitochondrial inner membrane (NADH and NADH+Oligo for S1). These results, together with (i) ATP synthesis and hydrolysis rates, which are primarily affected in the S1 strain (Supplemental Figure 1A), (ii) normal levels and assembly of respiratory complexes observed via BN-PAGE in all strains (Supplemental Figure 1B), (iii) preservation of mitochondrial membrane potential as shown by rhodamine 123 fluorescence quenching, though partially uncoupled in S1 (Supplemental Figure 1C), and (iv) normal synthesis of mitochondrial DNA-encoded proteins across all strains (Supplemental Figure 1D), indicate that the mitochondria in the three strains exhibit comparable OXPHOS content and activity. Interestingly, during mitochondrial sample preparation, we consistently observed that the S1 strain, in which mitochondria are partially uncoupled, yielded a higher mitochondrial content per cell. This observation suggested an overall increase in mitochondrial abundance (see Materials and Methods for details on mitochondrial preparation). To validate this, we performed a similar analysis on whole cells (Figure 1B, Cells), measuring oxygen consumption using ethanol as an electron donor under different conditions: in the presence of the membrane potential uncoupler CCCP (Ethanol+CCCP, maximal respiration), under phosphorylating conditions (Ethanol, state 3), and with triethyltin, an ATP synthase inhibitor (Ethanol+TET, state 4). Under all conditions, the S1 strain displayed a 30–50% increase in respiratory parameters relative to YE1 and S1+ε, potentially indicating an approximate doubling of mitochondrial content per cell.

To further investigate this phenomenon, we quantified cytochrome and mitochondrial protein content in both whole cells and isolated mitochondria (Figure 1C). In whole cells, the S1 strain exhibited a 20–30% increase in cytochrome levels (Figure 1C, Cells), an effect not observed in isolated mitochondria (Figure 1C, Mitochondria). Supporting this, immunoblot analysis of various mitochondrial proteins revealed a 30–50% increase in ATP synthase subunits in whole cells relative to isolated mitochondria (Figure 1D, Cells/Mitochondria), reinforcing the findings from cytochrome quantification. Importantly, these variations were largely reversible, as most respiratory activities, cytochrome levels, and protein abundances in S1+ε strain returned to those observed in the YE1 strain, with the notable exception of porin (Figure 1B-D).

Unexpectedly, in the rescued S1+ε strain, cytochrome levels in whole cells were reduced by approximately 30–40% compared to the wild-type YE1 strain, a result inconsistent with the respiratory activity measurements (Figure 1B/1C, Cells). A slight decrease in both respiratory activity and cytochrome content was also evident in isolated mitochondria, suggesting a modest reduction in overall enzyme levels and/or potentially mitochondrial content in these cells.

To broaden the scope of our analysis, we also examined the cytosolic protein phosphoglycerate kinase (Pgk), which showed increased abundance in the S1 strain relative to YE1 and S1+ε. This observation suggests that the cellular alterations in the S1 strain may extend beyond mitochondria, potentially reflecting more global adaptations (see below).

#### 2.1.2. Influence of the mitochondrial uncoupling on mitochondrial structure

Given that the S1 mutant shows partial uncoupling and increased mitochondrial content, we aimed to investigate how these changes impact the mitochondrial network. We transformed the three yeast strains (YE1, S1 and S1+ε) with a plasmid that constitutively expresses GFP targeted to the mitochondria (Supplemental Table 1) and used fluorescence microscopy to visualize the mitochondrial network in cells grown in a respiratory medium (glycerol/ethanol). Only the S1 strain displayed significant alterations, with its mitochondrial network fragmenting into small spheres (Figure 2A). Electron microscopy further revealed no changes in the mitochondria shape, size, density across the three strains and cristae structure (Figure 2B and Supplemental Figure 2A), however, there was a significant increase in their number in the S1 strain (Figure 2B and Supplemental Figure 2B). Thus, the primary characteristic of the S1 mutant is a fragmented mitochondria network and a substantial increase in mitochondrial content, though its mitochondria are otherwise similar to those in the wild-type YE1 and the S1+ε rescue strains, consistent with previous findings.

**Figure 2 -.**
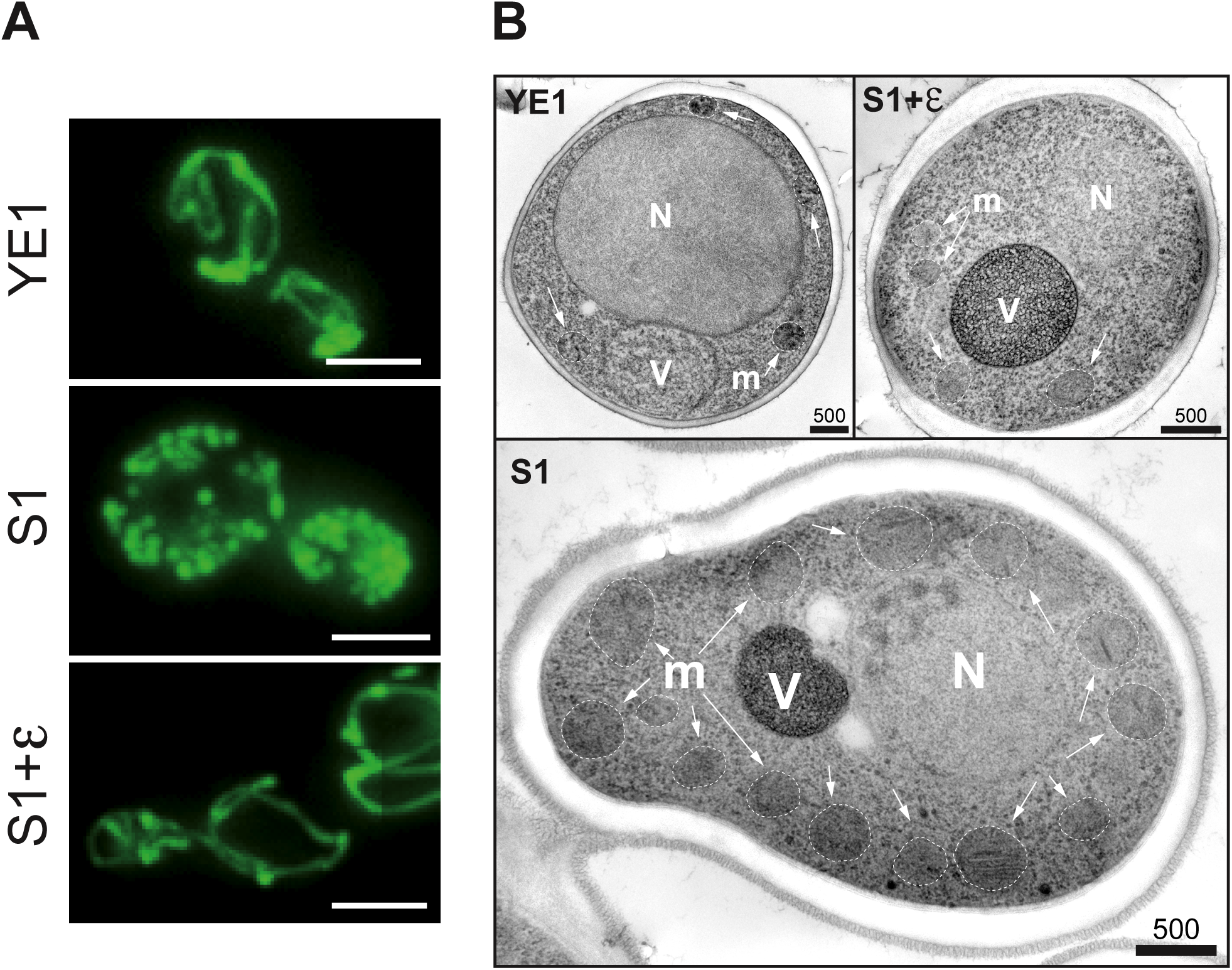
Mitochondrial structure. **(A)** Fluorescence microscopy of wholes cells expressing mitochondrial GFP protein. Scale bars: 5 µM. **(B)** Electron microscopy images of mitochondria per section (cross section corresponds to a cell where nucleus and vacuole were observed). The arrows indicate mitochondria (m) and the vacuole and nucleus were indicated respectively by V and N. Scale bars (nm). See also Supplemental Figure 2A/B.

#### 2.1.3. Influence of the mitochondrial uncoupling on mitochondrial biogenesis

Substantial increase in the number of mitochondria per cell in S1 mutant cells could potentially linked to enhanced mitochondrial biogenesis. To investigate this biogenesis, we employed two approaches.

First, we transformed the three strains with a reporter plasmid expressing β-galactosidase under the control of the SDH3 promoter (Succinate DeHydrogenase 3), a promoter regulated by Hap transcription factors involved in mitochondrial biogenesis during the diauxic transition [33]. In contrast to YE1 and S1+ε rescue strains, S1 mutant exhibited a sharp increase (approximately 4-fold) in SDH3 promoter activity when grown in respiratory conditions (glycerol/ethanol), indicating strong Hap transcription factor activity and a significant stimulation of mitochondrial biogenesis (Figure 3A). To confirm that the induction of β-galactosidase was dependent on uncoupling, we examined β-galactosidase activity in YE1 and S1+ε strains after adding doxycycline to the culture medium (glycerol/ethanol), which blocks ε-subunit expression. In YE1, a slight stimulation of β-galactosidase activity was observed in response to doxycycline concentration, reflecting an induction of mitochondrial biogenesis, but this quickly led to total uncoupling and cell death (Supplemental Figure 3A) [32; 34]. As expected, S1+ε strain incubated with doxycycline present a strong stimulation of β-galactosidase activity, similar to that observed in the S1 mutant (Supplemental Figure 3B).

**Figure 3 -.**
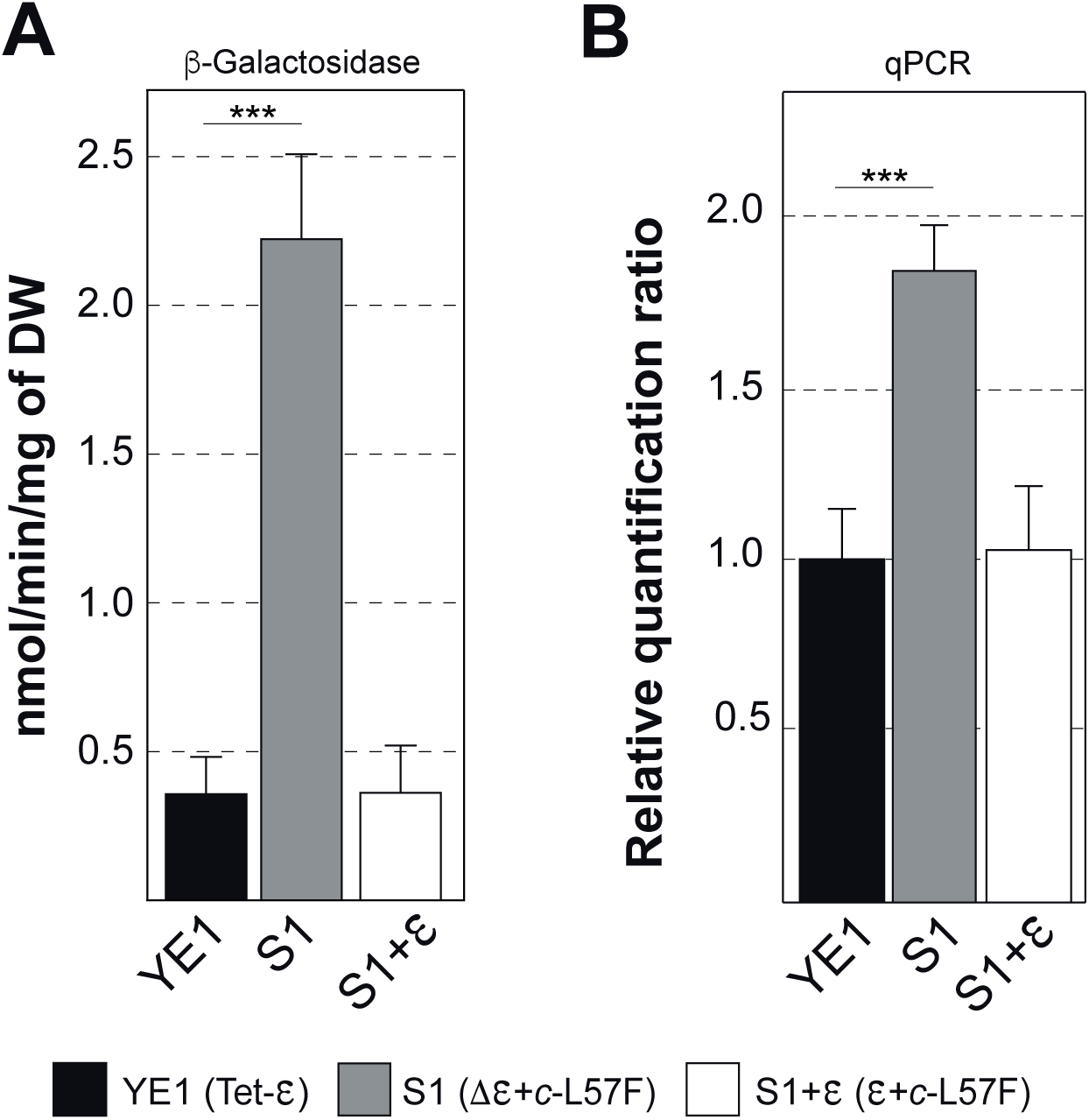
Mitochondrial proliferation. **(A)** Mitochondrial biogenesis was followed by measuring the activity of the transcription factors HAPs with a widely used ß-galactosidase reporter gene, (p161 plasmid, pSDH3-LacZ). DW, dry weight. **(B)** Relative quantification ratio of mitochondrial *ATP9* gene in comparison to the nuclear actin reference gene. See also Supplemental Figure 3A/B and Supplemental Table 2.

Second, we measured the amount of mitochondrial DNA (mtDNA) in the cells using qPCR (See Supplemental Table 2, Materials and Methods). An increase in mtDNA copy number is a typical marker of enhanced mitochondrial biogenesis [35]. We used the actin gene as a nuclear control and the ATP9 gene (c-subunit) of ATP synthase as the mitochondrial gene for qPCR experiments. The S1 mutant showed a large increase in mtDNA when grown in glycerol/ethanol compared to the wild-type YE1 and rescue S1+ε strains, indicating that mitochondrial biogenesis correlates with an accumulation of mtDNA (Figure 3B).

In conclusion, partial mitochondrial uncoupling appears to stimulate mitochondrial biogenesis, resulting in nearly double the number of mitochondria per cell in the S1 mutant, with these mitochondria being indistinguishable from those in wild-type cells. Additionally, this stimulation seems to be fully reversible, as observed in the rescue S1+ε cells when mitochondrial uncoupling is reversed by re-expressing the ε-subunit.

### 2.2. Mitochondrial uncoupling leads to a metabolism remodeling

#### 2.2.1. Influence of the mitochondrial uncoupling on metabolism

A notable finding from the biochemical analyses, particularly the immunoblot analysis of S1 mutant cells (Figure 1D, Cells), was the substantial increase in phosphoglycerate kinase (Pgk), a cytosolic glycolysis enzyme involved in the conversion of 1,3-BP-glycerate to 3P-glycerate. However, this enzyme is not regulated by Hap transcription factors, suggesting an additional adaptive response beyond mitochondrial biogenesis. To characterize this response, we performed a global and quantitative differential proteomic analysis (2D-DiGE) to compare protein expression in S1 mutant cells with wild-type YE1 cells.

The 2D-DiGE analysis was conducted on whole cell extracts from eight samples, four controls (YE1) and four mutants (S1). Each gel generated three images corresponding to Cy2, Cy3, and Cy5-labeled samples. These 12 images were analyzed using DeCyder software to detect proteins and calculate their average abundances. We considered changes with a 95% confidence interval (p<0.05) and standardized average spot volume ratios exceeding 1.5. This analysis identified 201 protein spots significantly altered in the S1 mutant, with 128 spots increased and 73 spots decreased compared to YE1 controls. These spots are shown on a representative 2D-DiGE gel image in Supplemental Figure 4A. Selected spots were digested in gel and analyzed by MALDI-TOF-TOF. A database search using peptide mass fingerprint spectra identified proteins in 183 of the 201 spots. Fifteen spots were unidentified, three had only one peptide and were considered unidentified, and 46 spots contained two or three different proteins. In total, 137 different spots corresponding to 78 different proteins were altered in the S1 mutant, detailed in Supplemental Table 3, including gene name, accession number, peptide number, peptide sequence, score, enrichment factor, theoretical molecular weight, pI values, and a brief description.

A subsequent database search categorized these proteins by function and predicted subcellular localization: 36% were cytosolic, 22% ribosomal, 10% associated with the nucleus, vacuole, or endoplasmic reticulum, and 32% mitochondrial (Supplemental Table 3, Figure 4A). The 78 altered proteins were divided into two major functional categories. The primary group (55%, with 86% up-regulated) is involved in metabolic pathways, mainly energy metabolism (84% glycolysis and TCA cycle, 16% stress responses). The second group (22%, exclusively down-regulated) is associated with the translation machinery. Remaining proteins, mostly down-regulated, are involved in protein degradation (6%), chaperones (6%), or cell homeostasis regulation (11%). Figure 4B provides an overview of these proteins and their metabolic pathways.

**Figure 4 -.**
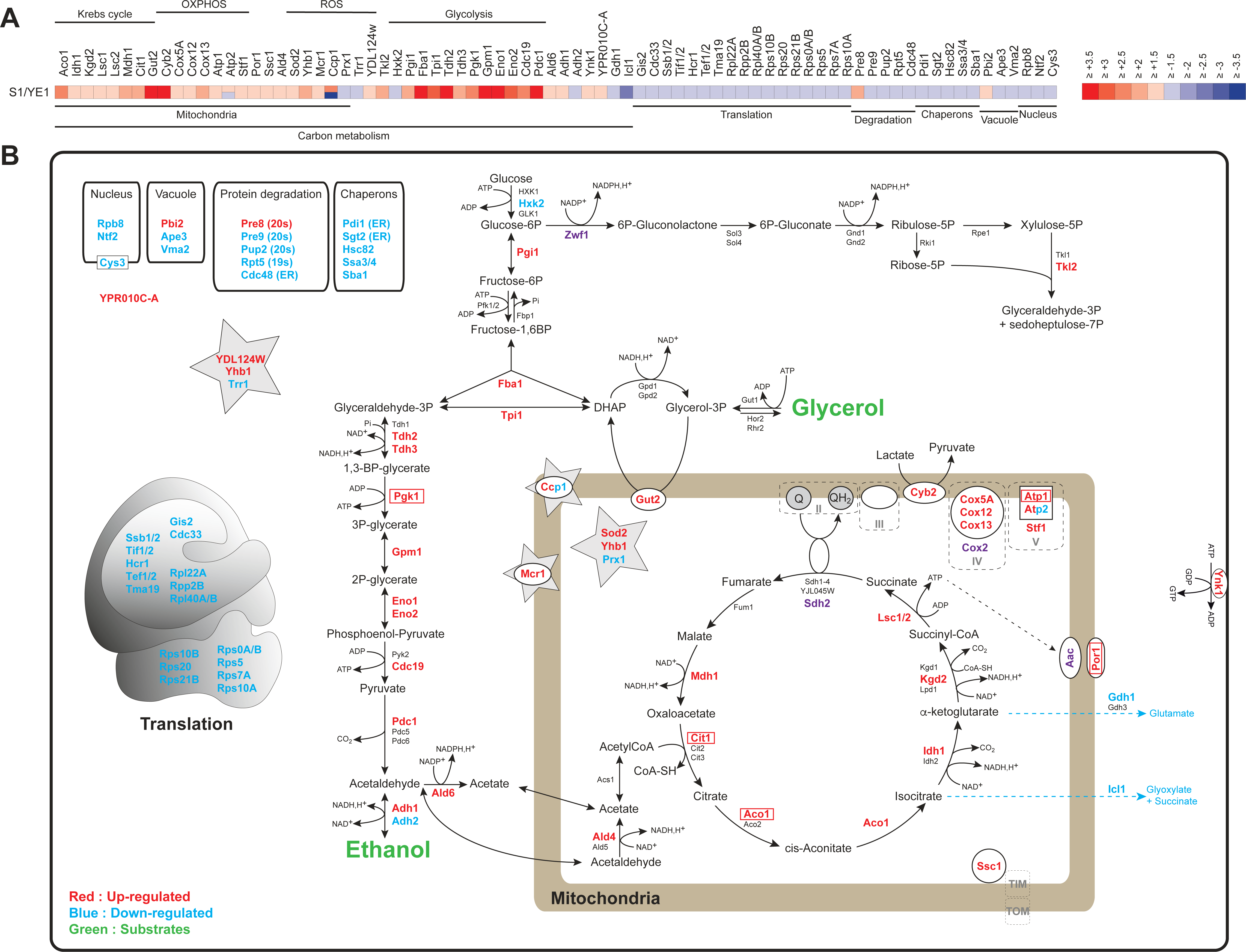
Biological pathways up or down-regulated in the S1 mutant strain. **(A)** List of 78 quantified proteins and heat map representation of up-regulated or down-regulated proteins (S1/YE1 > 1.5). See also Supplemental Table 3 and Supplemental Figure 4A/B/C/D). **(B)** Overview of up or down-regulated proteins. Substrates were indicated in green, up-regulated and down-regulated proteins in red and blue respectively. The names of enzymatic isoforms or subunits complexes are written in black. Proteins involved in ROS detoxification were indicated by a grey star (Yhb1, Trr1, Sod2, Prx1, Mcr1, CCP1 and YDL124W). Proteins up-regulated and framed were verified by immuno-blotting or by measuring their activity (Por1, Atp1, Ccp1, Pgk1, Cit1 and Aco1). Protein indicated in purple (Zwf1, Cox2, Sdh2 and Aac) have been only quantified by immunoblot (See also Supplemental Figure 1D and Supplemental Figure 4C/D).

To validate these findings, we monitored the expression levels of several altered proteins and other cytosolic and mitochondrial proteins using immunoblotting and enzymatic activity assays. This confirmed increased levels of Pgk, porin (Por1), mitochondrial ATP synthase subunit 1 (Atp1), cytochrome-c peroxidase (Ccp1), aconitase (Aco1), and mitochondrial citrate synthase (Cit1) (Figure 1D and Supplemental Figure 4B-D). Additionally, we tested other proteins not identified by 2D-DiGE with available antibodies (Cox2, Sdh2, Atp11, Zwf1, Hsp60, and Ade13). Only Cox2, Sdh2, Aac, and Zwf1 levels increased in the S1 mutant, while Atp11, Hsp60, and Ade13 levels remained unchanged, indicating specific metabolic alterations (Figure 4B, in purple and Supplemental Figure 4C-D). Our 2D-DiGE analysis revealed a significant increase in enzymes involved in glycerol and ethanol metabolism, particularly glycolysis, the TCA cycle, and some OXPHOS complexes. These observations are in agreement with our previous results and suggest that ATP production in S1 cells predominantly occurs via substrate-level phosphorylation mediated by succinyl-CoA ligase (Lsc1/2), rather than through the non-functional ATP synthase. Supporting this, alternative TCA cycle pathways were down-regulated, including isocitrate lyase (Icl1) for the glyoxylate cycle and glutamate dehydrogenase (Gdh1) for glutamate formation (Figure 4B).

Interestingly, chaperones, proteasome components, and translation machinery proteins, which are energy-consuming, were reduced. This reduction suggests a cellular response to low mitochondrial ATP production and could explain the slow growth of S1 mutant cells. Additionally, several proteins involved in reactive oxygen species (ROS) metabolism exhibited altered expression levels. Mitochondrial superoxide dismutase (Sod2), cytosolic and mitochondrial nitric oxide oxidoreductase (Yhb1), mitochondrial NADH-cytochrome b5 reductase (Mcr1), mitochondrial cytochrome-c peroxidase (Ccp1), and NADPH-dependent alpha-keto amide reductase (YDL124W) were upregulated. In contrast, mitochondrial peroxiredoxin (Prx1) and cytoplasmic thioredoxin reductase (Trr1) were downregulated. These changes suggest that metabolic adjustments for energy production may influence or result in an increased ROS generation.

#### 2.2.2. Impact of metabolic remodeling on ROS production

Given the increase in proteins involved in ROS detoxification, we measured ROS production in whole cells using the fluorescent probe dihydroethidium. We observed a significant increase in cellular ROS production in S1 cells, which exhibit both stimulated mitochondrial biogenesis and adapted energy metabolism compared to the wild-type YE1 and rescue S1+ε strains (Figure 5). Additionally, the growth of S1 mutants slightly improved when ROS scavengers, such as N-acetyl cysteine (NAC) and ascorbic acid (ASC), were added to the culture media (glycerol/ethanol) (Supplemental Figure 5), suggesting that ROS may contribute to the slower growth of S1 cells compared to wild-type cells.

**Figure 5 -.**
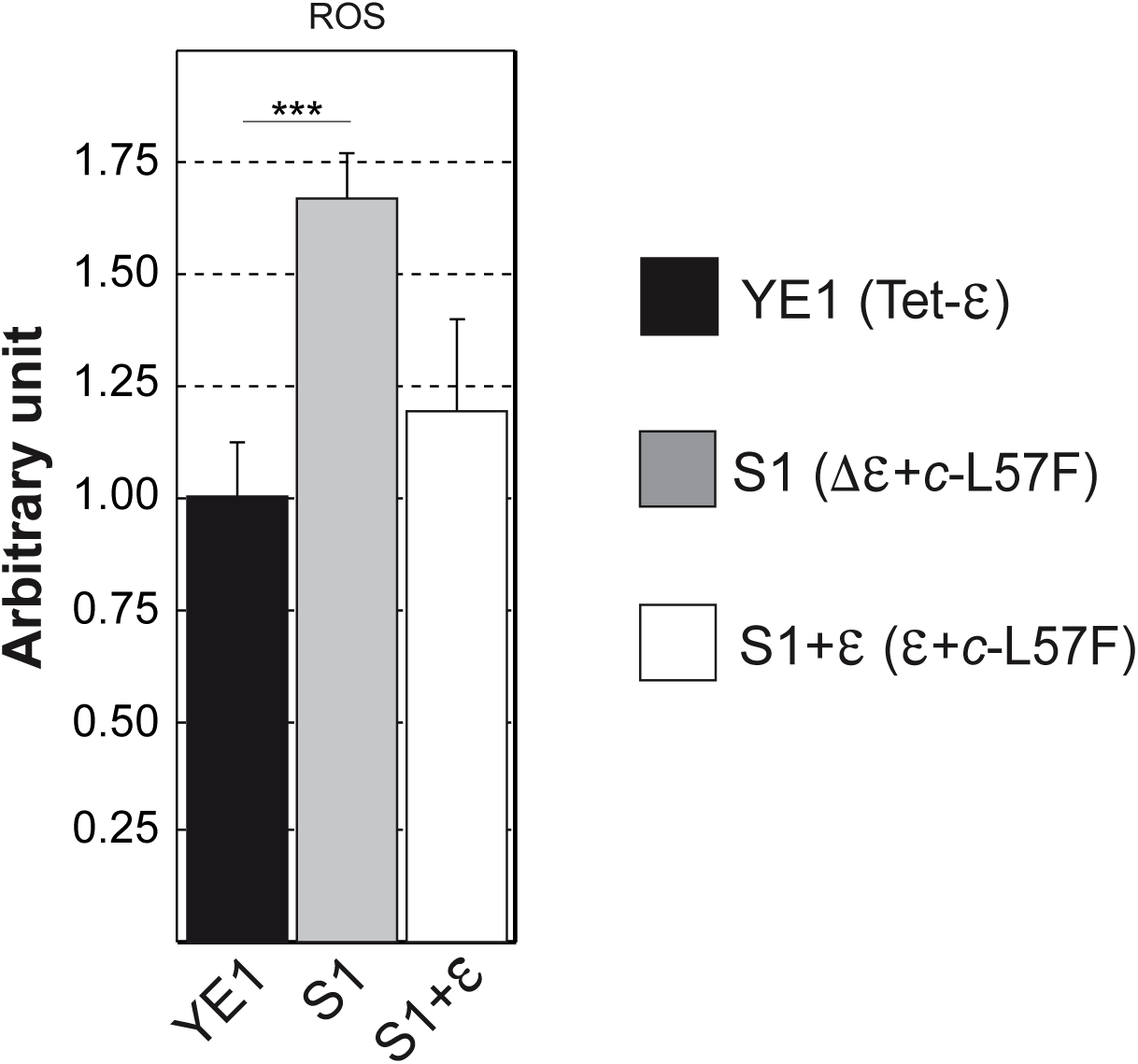
Rate of ROS production. The rate was normalized against ROS production of the YE1 strain. YE1 (Tet-ε), S1 (Δε+*c*-L57F) and S1+ε (ε+*c*-L57F) strains were grown on glycerol/ethanol medium at 28°C. Data are represented as mean ± SEM of at least three independent cell cultures. See also Supplemental Figure 5.

#### 2.2.3. Impact of the succinyl-CoA ligase on S1 energy metabolism

We hypothesized that the metabolic adaptation in S1 mutant cells allows for ATP production via substrate phosphorylation by the succinyl-CoA ligase enzyme (Lsc1/2, EC 6.2.1.4). This enzyme catalyzes the nucleotide-dependent conversion of succinyl-CoA to succinate, producing one ATP molecule, and consists of α (Lsc1) and β (Lsc2) subunits, both necessary for its catalytic activity. In S1 cells, both subunits are up-regulated (Figure 4B and Supplemental Table 3). Consequently, deleting this enzyme should be lethal for S1 cells grown in glycerol/ethanol conditions. Before deleting *LSC1* gene in S1 mutant cells, we needed to confirm that this activity is non-essential for wild-type cell growth under respiratory conditions. Deleting the *LSC1* gene in wild-type W303-1B showed no impact on both fermentation and respiratory growth (Figure 6), which is consistent with previous studies [36]. To avoid significant effects on cell growth (lethality or growth limitation) from *LSC1* deletion in S1 cells, we conducted the experiment in the rescue S1+ε strain. This strain allows specific inhibition of ε-subunit expression, leading to the S1 phenotype in the presence of doxycycline [32], thereby inducing metabolic alterations (as discussed earlier and shown in Supplemental Figure 3B). Under these conditions, and only in the presence of doxycycline, respiratory growth arrest was observed for S1+ε+Lsc1Δ cells (Figure 6), confirming our hypothesis. Additionally, deleting the other succinyl-CoA ligase subunit (Lsc2) in this context yielded the same results (Supplemental Figure 6), highlighting the crucial role of this enzyme in the viability of S1 mutant cells.

**Figure 6 -.**
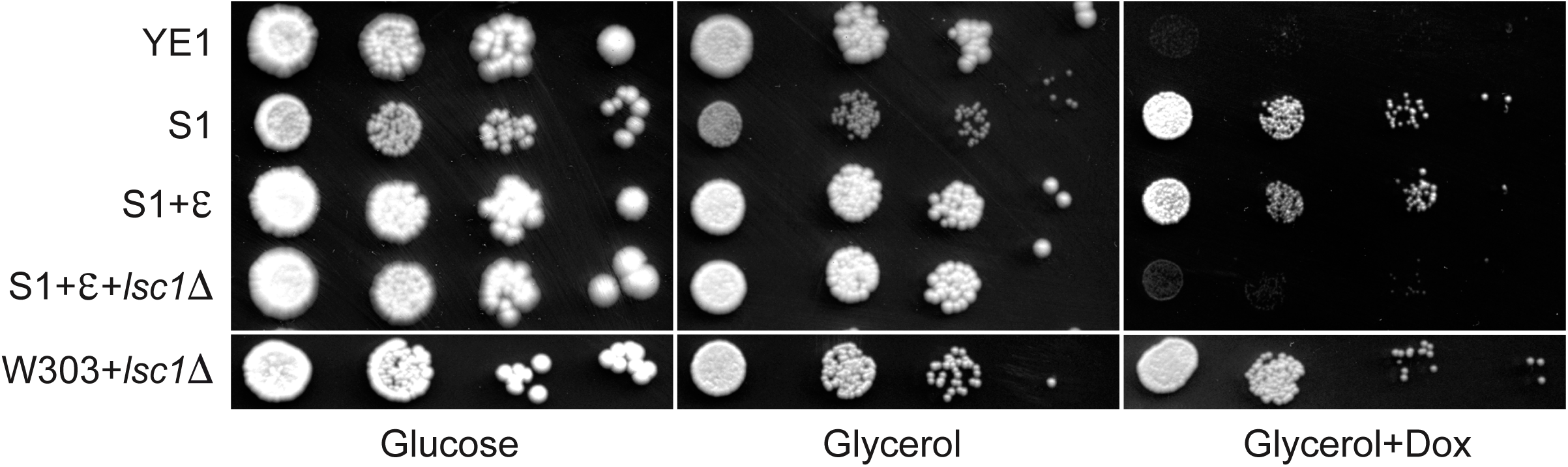
Participation in S1 mutant cells of succinyl-CoA ligase in ATP production. Drop tests of YE1 (Tet-ε), S1 (Δε+*c*-L57F), S1+ε (ε+*c*-L57F), S1+ε *lsc1Δ* and W303-1B+*lsc1Δ* strains on glucose, glycerol and glycerol+doxycycline (10µg/ml) media and grown for 3–5 days at 28°C. See also Supplemental Figure 6.

### 2.3. Mitochondrial biogenesis and metabolic remodeling are related to mitochondrial proton-motive force adjustment

Multiple signals or combinations of signals, likely originating from the mitochondria, could be responsible for these significant alterations. As a reminder, the S1 mutant cells have deficient ATP synthase, partially uncoupled mitochondria with low mitochondrial potential, and high ROS accumulation. Therefore, we focused on variations in mitochondrial potential, proton-motive force (PMF), and ROS production. Although ROS is typically described as lowering mitochondrial biogenesis in yeast, its role is still debated in mammalian cells [30]. To explore this, we investigated the influence of molecules that modify mitochondrial potential and/or PMF and their effects on mitochondrial biogenesis and ROS production using our β-galactosidase reporter system and the fluorescent probe dihydroethidium respectively. Additionally, to avoid the specific context of the S1 mutant, we conducted these experiments on a wild-type yeast strain (W303-1B), bypassing potential influences from the c-L57F ATP synthase mutation.

Initially, we analyzed β-galactosidase activity induction in WT yeast treated with the protonophore CCCP to mimic mitochondrial uncoupling observed in the S1 mutant. Under these conditions, a slight stimulation of β-galactosidase activity was observed at low CCCP concentrations (considerably less than in the S1 mutant). At higher CCCP concentrations, there was no stimulation of β-galactosidase activity due to major cell membrane alterations related to total uncoupling, which is non-specific to mitochondria and leads to cell death (Supplemental Figure 7A). These conditions are likely too detrimental for the cells; therefore, we used other uncoupling molecules (valinomycin and nigericin) with more selective action on mitochondrial membranes in yeast [37]. Valinomycin mainly affects mitochondrial potential through potassium exchange without influencing PMF, whereas nigericin exchanges potassium for protons, maintaining mitochondrial potential but collapsing PMF. We measured both β-galactosidase activity and ROS production in cells cultured with each molecule under conditions that maintained cell growth. A strong stimulation of β-galactosidase activity was observed in cells treated with increasing nigericin concentrations, in contrast to valinomycin; higher concentrations of valinomycin resulted in cell death (Figure 7A). We also evaluated additional inhibitors of the respiratory chain (antimycin, myxothiazol), mitochondrial ATP synthase (oligomycin), and another protonophore (2,4-dinitrophenol, DNP). A small stimulation was observed with low DNP concentrations, similar to CCCP (Supplemental Figure 7A/7B). Unexpectedly, antimycin inhibited β-galactosidase expression, unlike myxothiazol, and never stimulated β-galactosidase activity (Supplemental Figure 7B). The mechanism underlying the inhibition by antimycin remains unclear at this stage, and further studies will be necessary to elucidate it.

**Figure 7 -.**
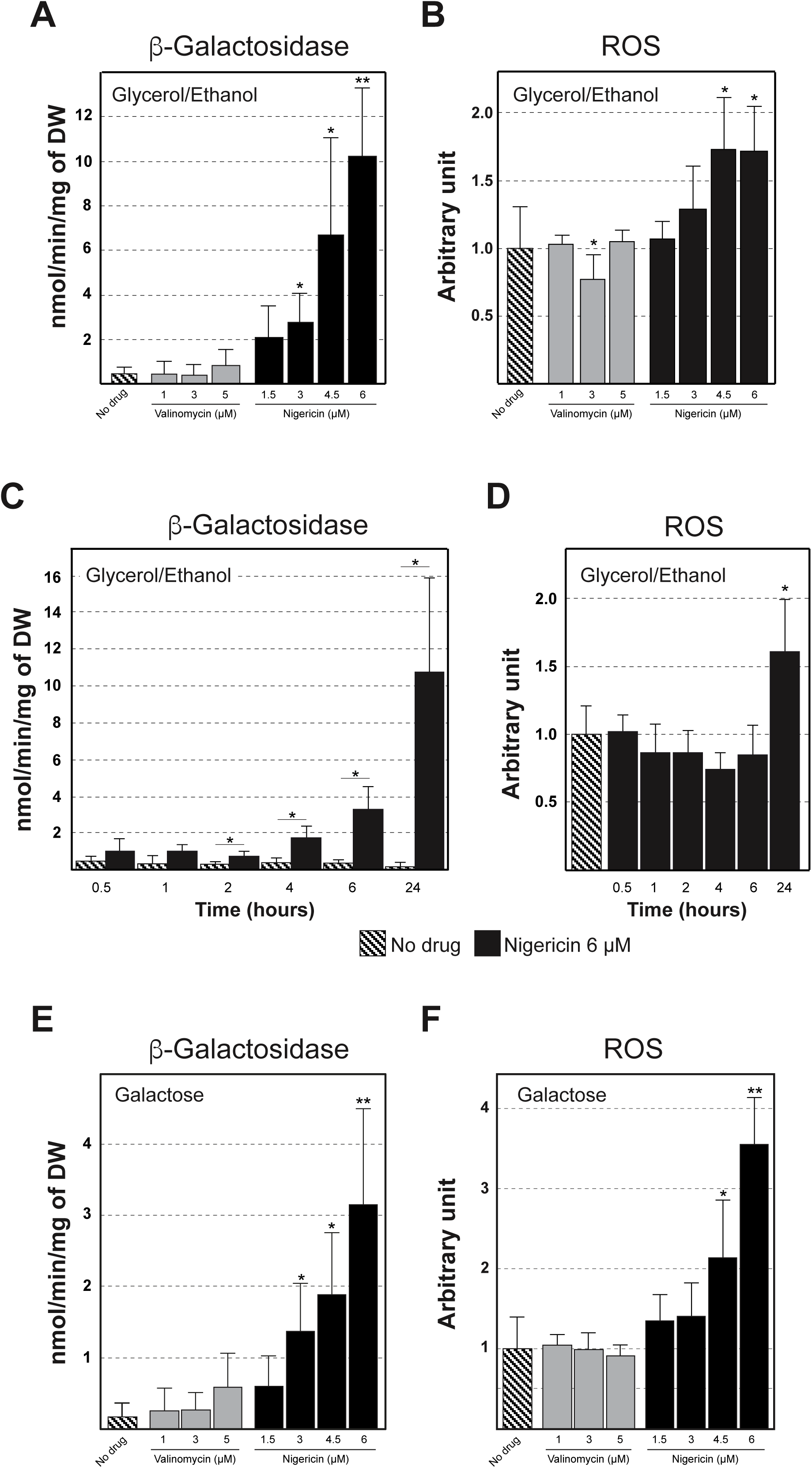
Induction of mitochondrial biogenesis in wild-type cells. **(A)** Mitochondrial biogenesis was followed by measuring the activity of the ß-galactosidase reporter gene, (pSDH3-LacZ). Wild type W303-1B+pSDH3 cells were grown in synthetic 2% glycerol/ethanol medium in absence/presence of valinomycin or nigericin (4 to 5 generations, OD=2) at 28°C. **(B)** The rate of ROS production in cells was normalized against ROS production of the W303-1B+pSDH3 strain grown in 2% glycerol/ethanol medium without addition of valinomycin or nigericin. Cells were grown as previously described in A. **(C)** Monitoring mitochondrial biogenesis induction in function of time after addition of nigericin by measuring the activity of the ß-galactosidase reporter gene, (pSDH3-LacZ). Cells were grown in 2% glycerol/ethanol at 28°C. **(D)** Rate of ROS production in function of time after addition of nigericin. ROS production was normalized against ROS production of the W303-1B+pSDH3 strain grown in 2% glycerol/ethanol medium without addition of nigericin at 28°C. **(E/F)** Exactly as described in A/B but this time galactose was used instead of glycerol/ethanol as substrate. Data are represented as mean ± SEM of at least three independent cell cultures. See also Supplemental Figure 7.

Similarly, a large increase in ROS production was observed with nigericin, in contrast to valinomycin, antimycin, myxothiazol, oligomycin, and DNP (Figure 7B and Supplemental 7C), suggesting that mitochondrial metabolism and biogenesis adaptations are ROS producers. However, it is unclear whether PMF or ROS is altered first. To distinguish between ROS and PMF, we conducted a kinetic study on β-galactosidase activity induction and ROS production in cells cultured with 6 µM nigericin, a concentration that induces pronounced reporter activity stimulation (Figure 7A). After 4 to 6 hours of nigericin treatment, an increase in β-galactosidase activity was noted, consistent with the beginning of mitochondrial biogenesis stimulation (Figure 7C), while ROS levels remained constant or slightly decreased (Figure 7D). High ROS production was observed only after 24 hours, likely due to metabolic modifications (Figure 7D).

Finally, we investigated whether this response was similar in fermentative conditions using galactose as a carbon source, where ATP production mainly relies on glycolysis. We measured β-galactosidase activity stimulation and ROS production in wild-type yeast grown under these conditions with valinomycin or nigericin. Again, a strong stimulation of β-galactosidase activity and ROS production was observed only with nigericin, suggesting that the carbon source is not related to triggering these adaptation mechanisms (Figure 7E/F).

In conclusion, changes in PMF trigger robust mitochondrial biogenesis and profound metabolic remodeling in cells, recapitulating the phenotype of S1 mutant cells and underscoring the central role of mitochondrial PMF regulation in adapting energy metabolism.

## 3. DISCUSSION

Controlling and regulating cellular homeostasis, particularly mitochondrial homeostasis, is critical for our cells. The mechanisms involved are numerous, complex, and not always clearly identified. Several studies have shown that mitochondrial metabolism adjusts to meet cellular energy demands, resulting in modulations of enzyme activities or variations in mitochondrial mass [38–43]. Moreover, mitochondrial accumulation has been observed during aging and in many pathologies, including mitochondrial diseases characterized by OXPHOS defects from mutations in mitochondrial or nuclear genes [31; 44–57]. However, the molecular mechanisms underlying these variations in mitochondrial mass remain poorly understood.

Our studies on ATP synthase biogenesis [32; 34] and yeast models of mitochondrial diseases [31] allowed us to isolate an ATP synthase mutant (S1) capable of surviving mitochondrial uncoupling, exhibiting a high stimulation of mitochondrial biogenesis and an altered mitochondrial network (fragmented versus tubular). Interestingly, a similar situation occurs in mammalian cells, particularly in brown adipose tissue, where cells express the mitochondrial uncoupling protein “UCP1.” This protein uncouples mitochondria, leading to heat production and a significant decrease in mitochondrial ATP production [58]. Remarkably, these cells also have a large mitochondrial mass and a more fragmented mitochondrial network [59–63]. Some studies have also described that mitochondrial uncoupling can induce the transcription factor PGC1, which stimulates mitochondrial biogenesis [62;64; 65]. In the S1 yeast mutant, we observed both a strong increase in mitochondrial mass and a fragmented mitochondrial network, mirroring observations in brown adipose tissue (Figure 2 and 3). The decrease in mitochondrial potential and ATP production in the S1 mutant could be driving the fragmentation of the mitochondrial network, favoring fission over fusion. Indeed, ATP plays a crucial role in outer membrane fusion processes, while mitochondrial potential is important for inner membrane fusion [66].

We demonstrated that mitochondrial biogenesis in the S1 mutant is related to mitochondrial uncoupling (Figure 1) and controlled by Hap transcription factors (Figure 3). Whole-cell proteomic analysis in the S1 mutant revealed specific metabolic alterations aimed at compensating for low ATP production by ATP synthase through mitochondrial substrate phosphorylation (Figure 4 and 6), a stimulation of glycolysis, and a high ROS production (Figure 4, 5 and 7). Additionally, energy-consuming cellular mechanisms, such as translation and degradation, are diminished, reflecting the low growth rate of the S1 mutant compared to the wild-type strain (Figure 1, [32]). Interestingly, the study, expressing human UCP1 in yeast using a differential proteomics approach, also revealed an adaptation of mitochondrial content and an increased ROS production [67]. However, only the mito-proteome was analyzed, lacking a general view of cellular metabolism and compensatory pathways. Among the striking results of our proteomic analysis, and which were not observed in the previous study [67], was the overexpression of succinyl-CoA ligase subunits (Lsc1/2) allowing direct production of mitochondrial ATP unlike other succinyl-CoA ligases of other organisms that mainly use GTP [68]. Additionally, enzymes that shunt the Krebs cycle (Icl1 and Gdh1) were down-regulated, optimizing the Krebs cycle and promoting ATP production via succinyl-CoA ligase. Icl1 and Gdh1 are typically induced in respiratory conditions (ethanol) [69; 70] whereas here they were down-regulated suggesting accurate and specific adaptations of cellular metabolism in response to mitochondrial deficiency. It was also observed that unlike other glycolysis enzymes, Pfk and Fbp remained unchanged in the S1 mutant. This can be attributed to their activation mechanism, which involves phosphorylation and dephosphorylation processes, and therefore does not require an adjustment of their quantity [71–73].

We also observed an induction of enzymes involved in ROS detoxification (Sod2, Yhb1), reflecting an exacerbated metabolism which could come from the up-regulated mitochondrial Glycerol-3-dehydrogenase (Gut2) (Figure 4) [74; 75]. However, two enzymes (Trr1 and Prx1) are down-regulated whereas these are usually induced in respiratory conditions or in response to oxidative stress, down-regulation that could explain also the increased of ROS [76; 77]. The reason of this down-regulation is still unknown, but we can however note that they both use thioredoxin as a substrate, unlike the other up-regulated enzymes, suggesting that these detoxification enzymes are induced under conditions of oxidative stress that are not related to adaptations of energetic metabolism. Finally, we were able to show that the increased ROS production in the S1 mutant does not contribute to mitochondrial biogenesis (Figure 7), contrary to what is observed in some mammalian cells [78–80]. Indeed, cancer cells have inherently elevated ROS, presumably due to their high metabolic activity, which leads to up-regulation of the Nrf2 pathway [81; 82].

These findings strongly indicate that cellular metabolism adaptations are targeted, specific, and responsive to environmental conditions. Understanding the molecular mechanisms that coordinate these processes is critical for developing new strategies to understand and treat many human pathologies.

## MATERIALS AND METHODS

### 3.1. Strains and media

*E. coli* NEB-5alpha strain (BioLabs) was used for the cloning and propagation of plasmids. The *Saccharomyces cerevisiae* strains used, and their genotypes are listed in Table S1. The following rich media were used for the growth of yeast: 1% (w/v) yeast extract, 1% (w/v) peptone, 40 mg/l adenine, 2% (w/v) glucose or 2% (w/v) glycerol. The glycerol medium was buffered at pH 6.2 with 50 mM potassium phosphate, and 2% (w/v) ethanol was added after sterilization. We also used complete synthetic medium (CSM) (0.17% (w/v) yeast nitrogen base without amino acids and ammonium sulfate, 0.5% (w/v) ammonium sulfate, 2% (w/v) glucose and 0.8% (w/v) of a mixture of amino acids and bases from Formedium (Norfolk, UK). The solid media contained 2% (w/v) agar.

### 3.2. Oxygen consumption

The oxygen consumption of cells (fresh cultured media, OD_600_=2) was measured polarographically at 28°C using a Clark oxygen electrode in a 1 ml thermostatically controlled chamber. Respiratory rates were determined from the slope of a plot of 0_2_ concentrations *versus* time. Cell oxygen consumption of isolated mitochondria was measured in the respiration buffer (0.65 M mannitol, 0.36 mM EGTA, 5 mM Tris-phosphate, 10 mM Tris-maleate, pH 6.8) as previously described [83]. Isolated mitochondria were prepared by the enzymatic method [84]. The protein amounts were determined by the Lowry [85] method in the presence of 5% SDS and BSA as a standard.

### 3.3. Cytochrome content determination

The cellular and mitochondrial content of c+c_1_, b, and a+a_3_ hemes were measured as described in [86]. Briefly, cells were harvested in the same conditions used for polarography experiments (OD_600_=2), washed twice with distilled water and concentrated to obtain 2 ml of a cell suspension of about 50 optical density units at 600 nm. They were placed in a dual beam spectrophotometer (Varian Cary 4000) and a differential spectrum (from 500 to 650 nm) was performed between 1 ml of oxidized cells (1 µl of H_2_O_2_ 30% w/v, Sigma) and 1 ml of reduced cells (few grains of dithionite). For mitochondrial cytochrome content 5 mg of mitochondria preparation was used and treated as previously described for the cells. Calculations of cytochrome c+c_1_, cytochrome b and a+a_3_ contents were performed using an extinction coefficient of 18 mM^−1^ cm^−1^ (550/540 nm) 19 mM^−1^ cm^−1^ (564/575 nm) and 24 mM^−1^ cm^−1^ (605/630) respectively [87]. Cytochrome content is given in pmol per mg dry weight or mg of mitochondrial proteins.

### 3.4. ATP synthesis measurements

For ATP synthesis rate measurements, mitochondria (0.15 mg/ml) were placed in a 1-ml thermostatically controlled chamber at 28°C in respiration buffer. The reaction was started by the addition of 4 mM NADH and 1 mM ADP and stopped by 3.5% perchloric acid and 12.5 mM EDTA. The samples were then neutralized to pH 6.5 by the addition of KOH and 0.3 M MOPS. ATP was quantified in a luciferin/luciferase assay (Perkin Elmer) with an LKB bioluminometer. The specific ATPase activity was measured at pH 8.4 by using a previously described procedure [88]. The activity of the F_1_F_0_-ATP synthase in ATP production was assessed by oligomycin addition (20 µg/mg of protein).

### 3.5. Native gel electrophoresis

BN-PAGE experiments were carried out as described previously [89]. Briefly, mitochondrial extracts solubilized with digitonin to a protein ratio of 2 g/g were separated in a 3-13% acrylamide continuous gradient gel (Invitrogen). After electrophoresis, the gel was either stained with Coomassie Blue or incubated in a solution of 5 mM ATP, 5 mM MgCl_2_, 0.05% lead acetate, 50 mM glycine-NaOH, pH 8.4 to detect the ATPase activity [90] or transferred to nitrocellulose membranes and analyzed by western blotting.

### 3.6. Transmembrane potential (Δψ) measurement

Variations in transmembrane potential (Δψ) were evaluated in respiration buffer (0.65 M mannitol, 0.36 mM EGTA, 5 mM Tris-phosphate, 10 mM Tris-maleate, pH 6.8) by measuring the fluorescence quenching of rhodamine 123 with a SAFAS (Monte Carlo, Monaco) fluorescence spectrophotometer [91].

### 3.7. Fluorescence microscopy

Strains were transformed with a plasmid containing a mitochondrial GFP (pFL39-GFP, marker Trp1). For cell imaging, yeast cells were grown at 28°C in synthetic glycerol media until an OD600 between 0.5 to 1. Cells were washed once with phosphate buffer saline. 2.5 µl of the culture was spotted onto a glass slide and imaged at room temperature. Cells were observed in a fully automated inverted microscope (200M; Carl Zeiss, Inc.) equipped with a MS-2000 stage (Applied Scientific Instrumentation), a xenon light source (Lambda LS 175 W; Sutter Instrument Co.), a 100 × 1.4 NA Plan-Apochromat objective. For cell GFP imaging, we used FITC (excitation, HQ487/25; emission, HQ535/40; beamsplitter, Q505lp). Images were acquired using a camera (CoolSnap HQ; Roper Scientific). The microscope, camera, and shutters (Uniblitz) were controlled by SlideBook software (4.1; Intelligent Imaging Innovations).

### 3.8. Relative q-PCR experiments

DNA extraction was performed as described previously [92] from yeast culture grown in glycerol/ethanol. All q-PCR reaction mixtures were performed with 5X FIREPOL® EvaGreen qPCR Mix Plus (SOLIS BIODYNE) and quantifications were performed in a CFX Connect thermocycler (BIORAD). Data were analyzed using the mathematical model for relative quantification in real-time PCR [93]. Actin was used as internal references. q-PCR results and primers used in this study were presented in Figure 3 and Supplemental Table 2. We used for PCR 5 ng of DNA, 200 nM of each primer, 1X of Mix EVAgreen (5X FIREPOL® EvaGreen qPCR Mix Plus, SOLIS BIODYNE) and water to a final volume of 20 µl. The PCR program used was as follows: Step 1 - 95°C for 15 min; Step 2 - 95°C for 15 sec; Step 3 - 55°C for 20 sec; Step 4 - 72°C for 20 sec; Step 5 – Repeat 40 times from step 2.

### 3.9. *In vivo* labeling of mitochondrial translation products

These experiments were performed as previously described [94]. Briefly, strains were grown to early exponential phase (10^7^ cells/mL) in 10 mL of glycerol/ethanol rich media. The cells were harvested by centrifugation and washed twice with a low sulfate medium (W0-AS, W0 without ammonium sulfate; [95]). Cells were resuspended in 0.5 mL of W0-AS containing 1% glycerol/ethanol, supplemented with the corresponding auxotrophic markers but without methionine and cystein and 1 mM of a freshly prepared solution of cycloheximide and incubated at 20°C for 5 min; then 50 µCi of [^35^S]methionine-[^35^S]cysteine Mix (1,000 Ci/mmol, Perkin Elmer) was added. The reaction was stopped after 20 min, by the addition of 75 µL of 1.85 M NaOH, 1 M β-mercaptoethanol, and 0.01 M phenylmethylsulfonyl fluoride. An equal volume of 50% trichloroacetic acid was added, and the mixture was centrifuged in a microcentrifuge for 5 min at 14,000 rpm. The pellet consisting of precipitated proteins was washed once with 0.5 M Tris pH 8.9 and once with water and resuspended in 50 µL of sample buffer (2% (v/v) SDS, 10% (v/v) glycerol, 2.5% (v/v) β-mercaptoethanol, 0.06 M Tris-HCl pH 6.8 and 0.002% (w/v) bromophenol blue). The proteins were then separated on a polyacrylamide gel [96] and the resulting 12 or 15% gel was dried and revealed by scanning the 1 hour exposed image plate with a Fuji Bioimager. Amounts of [^35^S]-methionine-containing proteins in each lane were quantified using the ImageQuant software.

### 3.10. Citrate synthase activity

Isolated mitochondria from YE1, S1 and S1+ε of cultures grown on glycerol/ethanol condition were used to measured citrate synthase (EC 4.1.3.7) activity. The activity was determined spectrophotometrically by monitoring at 412 nm the oxidation of coenzyme A by 5,5’-dithiobis-2-nitrobenzoic acid (DTNB) over time in the following buffer: 50mM Tris-HCl (pH 7.5), 0.1mM acetyl-CoA, 0.2mM DTNB, and 0.5mM oxaloacetate. The enzyme activity was calculated using an extinction coefficient of 13 600M^−1^.cm^−1^ at 412 nm for DTNB.

### 3.11. Western-blot analysis

Polyclonal antibodies raised against yeast ATP synthase were used at a dilution of 1:50,000 for subunits Atp1; 1:10,000 for subunits Atp3, Atp4, *a*-Atp6, Atp7, *c*-Atp9, Atp15 and Atp16; and 1:10,000 for Aac and 1:5,000 for Cytb. Monoclonal antibodies against yeast porin and Pgk (from Molecular Probes) were used at a dilution of 1:5,000. Nitrocellulose membranes were incubated with peroxidase-labeled antibodies at a 1:5,000 dilution (Promega), and the blot visualization was conducted with ECL reagent (Pierce).

### 3.12. Freezing and freeze substitution for ultrastructural studies (MET)

The yeast pellets were placed on the surface of a copper EM grid (400 mesh) coated with formvar. Each grid was very quickly submersed in liquid propane precooled and held at −180°C using liquid nitrogen. Loops were transferred in a pre-cooled solution of 4% osmium tetroxide in dry acetone in a 1.8 ml polypropylene vial at −82°C for 72h (cryosubstitution), warmed gradually to room temperature, followed by three washes in dry acetone. The samples were infiltred progressively with araldite (epoxy resin Fluka). Ultrathin sections were contrasted with lead citrate. Specimens were observed with a HITACHI 7650 (80 kV) electron microscope (Electronic Imaging Pole of Bordeaux Imaging Center).

### 3.13. Mitochondrial biogenesis

Mitochondrial biogenesis was following by the activity of the β-galactosidase reporter gene [97] under the succinate dehydrogenase promoter gene (SDH3). A standard permeabilization procedure was used as described [98]. Briefly, after a preincubation period of 30 min at 30°C in a solution of 60 mM Na_2_HPO_4_, 40 mM NaH_2_PO_4_, 10 mM KCl, 1 mM MgSO_4_,7H_2_O, 1mM dithiothreitol and 0.2% w/v Sarcosyl, 0.4 mg/ml o-nitrophenyl-β-d-galactopyranoside (Sigma) was added and the mixture was incubated at 30°C. The reaction was stopped by the addition of 1 M Na_2_CO_3_. The samples were centrifuged for 1 min at 15,000 × g, and the absorbance of the supernatant was read at 420 nm. Activity is given in nmol per minute per mg dry weight.

### 3.14. Construction of a *Δlsc1* yeast strain

W303-1B and S1+Epsilon strains were transformed with the deletion cassette of LSC1 obtained by PCR amplification using plasmid pFL39 containing the tryptophan marker as a template and a pair of primers for LSC1 (LSC1-5, 5’-cgatttcatagaaaatttcttttgcagaccatttattcttcagtttgataATGTCTGTTATTAATTTCACAGGT AGTTC-3’/LSC1-3, 5’-tagaagtcttcctcttttgaaagcagttcattctcttcctaatataaTCATTTCTTAGCATTTTTGACGAAAT TTGC-3’, upper case indicate the coding sequence or TRP1 gene) according to a previously described procedure [99]. The transformants were selected on synthetic glucose media (CSM minus Uracil, Arginine and Tryptophan for S1+Epsilon and CSM minus Tryptophan for W303-1B) and confirmed by PCR analysis.

### 3.15. Reactive Oxygen Species (ROS) measurement

The YE1, S1 and S1+ε strains were grown in the same conditions used for polarography experiments (OD_600_=2). The cells were resuspended in 1 ml of phosphate buffer saline (PBS) before incubation with dihydroethidium (DHE, 5 µg/ml, Sigma) and further incubated in the dark for 5 min at room temperature. To quantify the number of cells displaying high ROS levels, at least 100,000 cells were counted in a BD Accuri^TM^ C6 flow cytometer and processed using CFlow Plus software. ROS production was controlled by addition of H_2_O_2_ (5 µl of 30% w/v stock solution, Sigma) in each sample. ROS production is given in arbitrary units.

### 3.16. Protein extraction, labeling and proteomic analysis

YE1 and S1 yeast strains were grown on rich glycerol/ethanol medium (OD_600_=2). Four different cultures for each strain were prepared, and protein extraction was performed in labeling buffer containing 30 mM Tris, 2 M thiourea, 7 M urea, 4% (w/v) 3-[3-(Cholamidopropyl)dimethylammonio]-1-propanesulfonate (CHAPS). The protein concentration was determined using the 2D Quant Kit (Amersham Biosciences, Uppsala, Sweden) and BSA as a standard. After extraction, proteins were used for a multiplexed analysis by two-dimensional difference in gel electrophoresis (DiGE) as described by Skynner et al. [100]. Protein labeling, bidimensional electrophoresis and image capture and analysis were exactly performed as described Lasserre et al. [101]. Spots having an abundance variation of at least 1.5-fold, *p*<0.05 were located on a gel (Supplemental Figure 4A).

### 3.17. Statistical analysis

Data are presented as mean ± SEM. All statistical analyses were performed using Student’s *t*-test (Two-tailed, unpaired), with \**p≤0.05*, \*\**p≤0.01* and \*\*\**p≤0.001*, respectively.

## ACKNOWLEDGMENTS

Scientific input is acknowledged from Anne Devin and Michel Rigoulet (IBGC, Bordeaux University) and Rodrigue Rossignol (MRGM, Bordeaux University). We particularly thank G. Lauquin for providing the immune serum against the Aac2 transporter, J. Velours and D. Brèthes for yeast ATP synthase antibodies, T. Langer for cytochrome *b* antibody, B. Daignan-Fornier for ß-galactosidase reporter plasmid (plasmid p161) and A. Sahin for the fluorescent-image acquisitions.

## AUTHOR CONTRIBUTIONS

Conceptualization, E.T., S.D-C. and J-P.d.R; Methodology, E.T.; Investigation, E.S., F.G., J-P.L., L.D., B.S., J.R. and E.T.; Resources, J.R. and B.S.; Writing - Original Draft, E.T., S.D-C. and J-P.d.R; Writing - Review & Editing, E.T., S.D-C. and J-P.d.R; Funding Acquisition, E.T. and J-P.d.R.; Supervision, E.T.

## DATA AVAILABILITY

Data will be made available on request.

## APPENDIX A. SUPPLEMENTARY DATA

Supplementary data to this article can be found online.

## DECLARATION OF GENERATIVE AI AND AI-ASSISTED TECHNOLOGIES IN THE WRITING PROCESS

During the preparation of this manuscript, the author(s) used AI for reformulation, as well as spelling and grammar correction. After using this tool/service, the author(s) reviewed and edited the content as needed and take(s) full responsibility for the content of the published article.

## Supplemental Information

**Supplemental Figure 1 (related to Figure 1) -.**
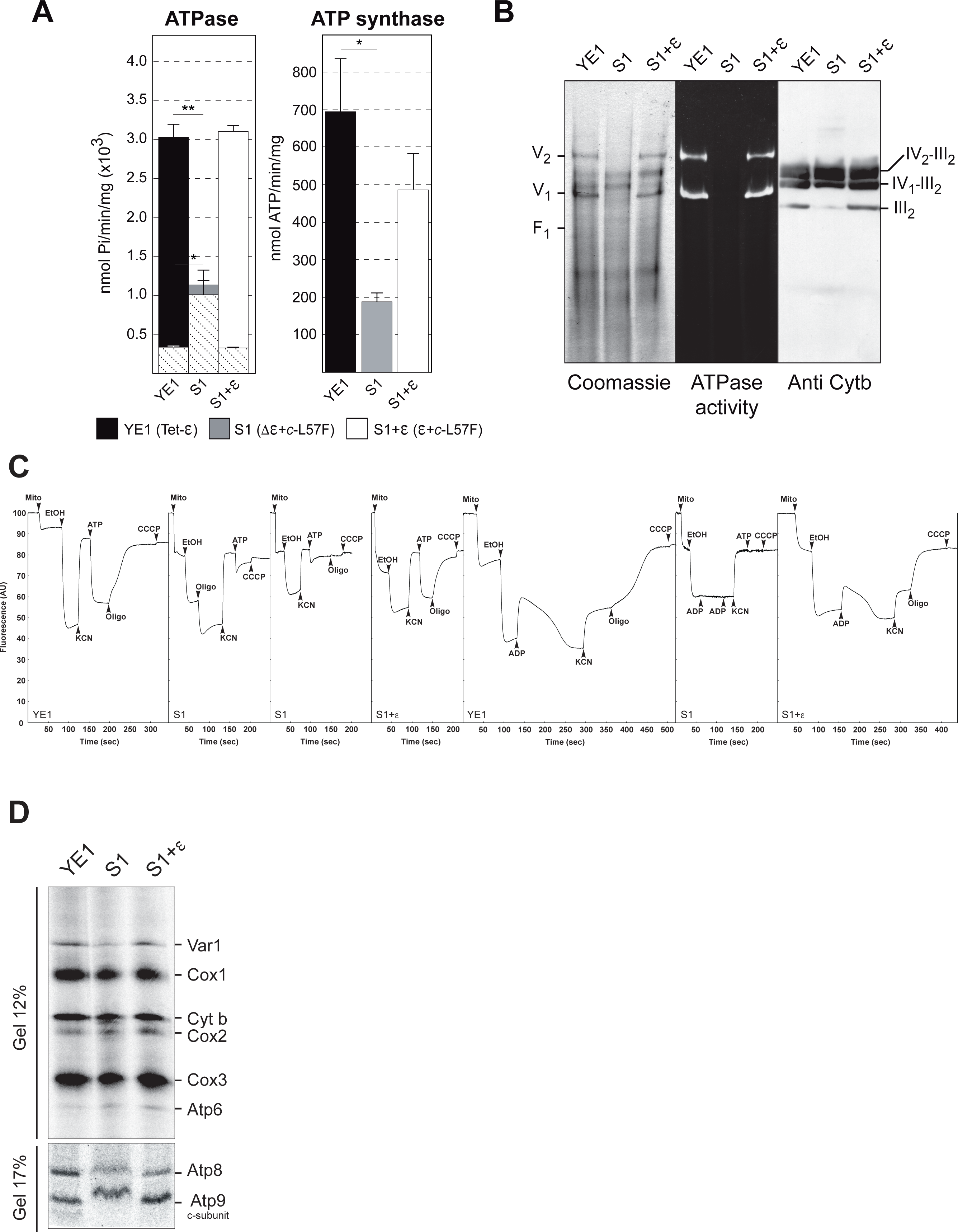
Mitochondria properties of the S1 mutant strain. **(A)** Measurements of ATPase/synthase activity on purified mitochondria from cells grown on glycerol/ethanol 2% at 28°C. The hatched bars representing the sensitivity to oligomycin. Data are represented as mean ± SEM of at least three inde-pendent cell cultures. **(B)** BN-PAGE analyses of the ATPsynthase. Mitochondria from YE1 (WT), S1 (Δε+c-L57F) and S1+ε (ε+c-L57F) mutants yeast strains from cells grown on glycerol/ethanol 2%, were solubilized with digitonin (2g digitonin/g of protein). After centri-fugation, the mitochondrial complexes were separated by BN-PAGE and the gels incubated with ATP and lead nitrate to reveal the ATPase activity (middle) followed by their staining with Coomassie Blue (left) or transfer onto a nitrocellulose membrane and probed with antibody against the Cytb (right). V1 and V2 correspond to monomeric and dimeric ATP synthase, respectively, and III2–IV2; III2–IV1 and III2 correspond to super-complexes III and IV. **(C)** Energization of the mitochondrial inner membrane. Variations in mitochondrial ΔΨ were monitored by the fluorescence quenching of rhodamine 123. The additions were 0.5 μg/ml Rhodamine 123, 0.15 mg/ml mitochondrial proteins (Mito), 10 μl of ethanol (EtOH), 2 mM potassium cyanide (KCN), 50 µM ADP, 4 μM CCCP, 0.2 mM ATP, and 4 μg/ml oligomycin (oligo). The fluorescence traces are representative of at least three experiments. **(D)** [35S] methionine pulse labeling of proteins translated in mitochondria (10,000 cpm per lane) shows relative variation of single-protein translation within each strain grown in glycerol/ethanol.

**Supplemental Figure 2 (related to Figure 2) -.**
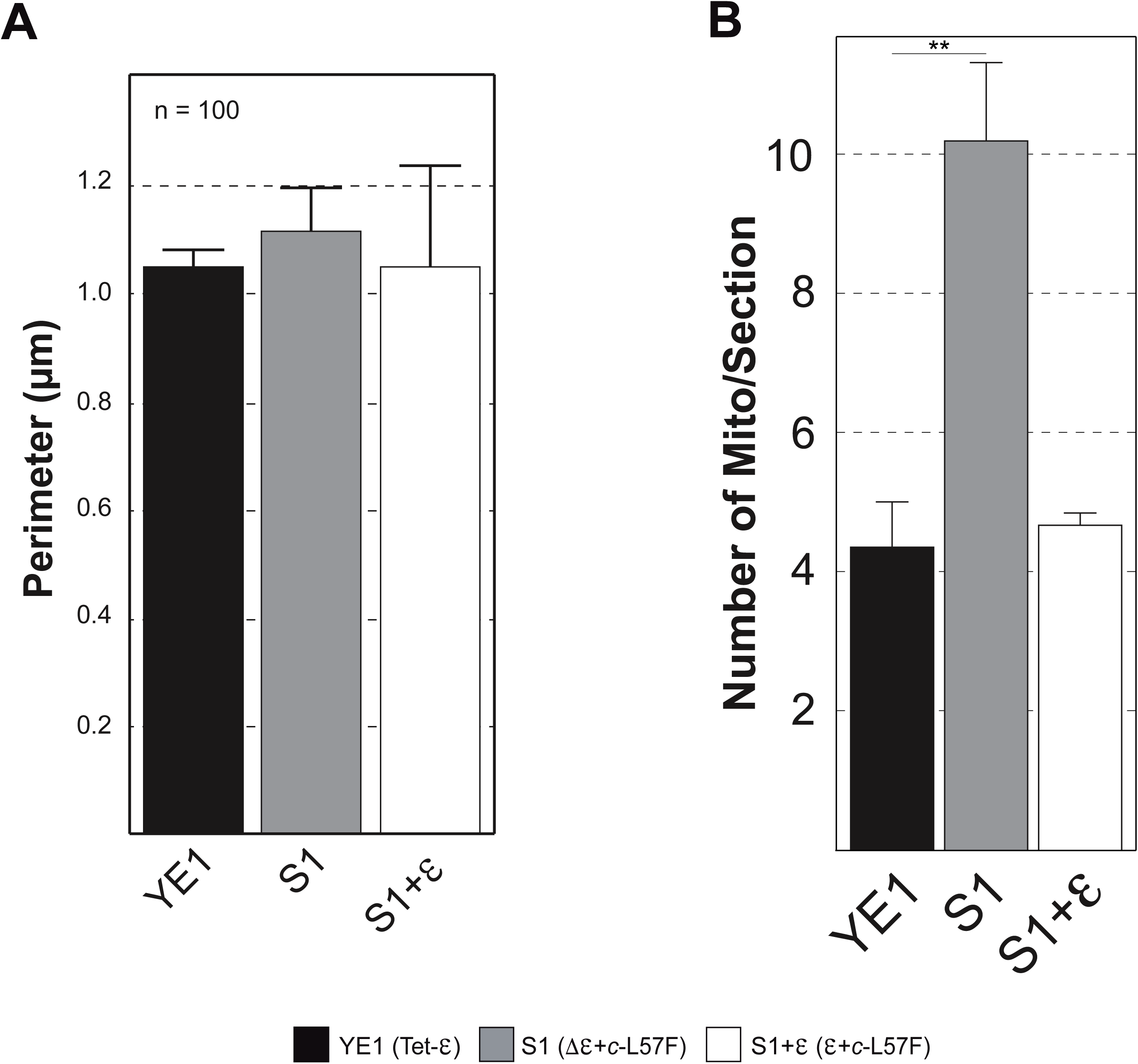
**(A) Ultrastructural analysis of mitochondria.** The perimeter of mitochondria was measured on a hundred mitochondrial electron micrographs using ImageJ software. Cells are grown on glycerol/ethanol medium. Data are represented as mean ± SEM of at least three independent cell cultures. **(B) Ultrastructural analysis of mitochondria**. Related quantifications of electron microscopy images of mitochondria per section (cross section correspond to a cell where nucleus and vacuole were observed). Cells are grown on glycerol/ethanol medium. Data are represented as mean ± SEM of at least 100 cells counted.

**Supplemental Figure 3 (related to Figure 3) -.**
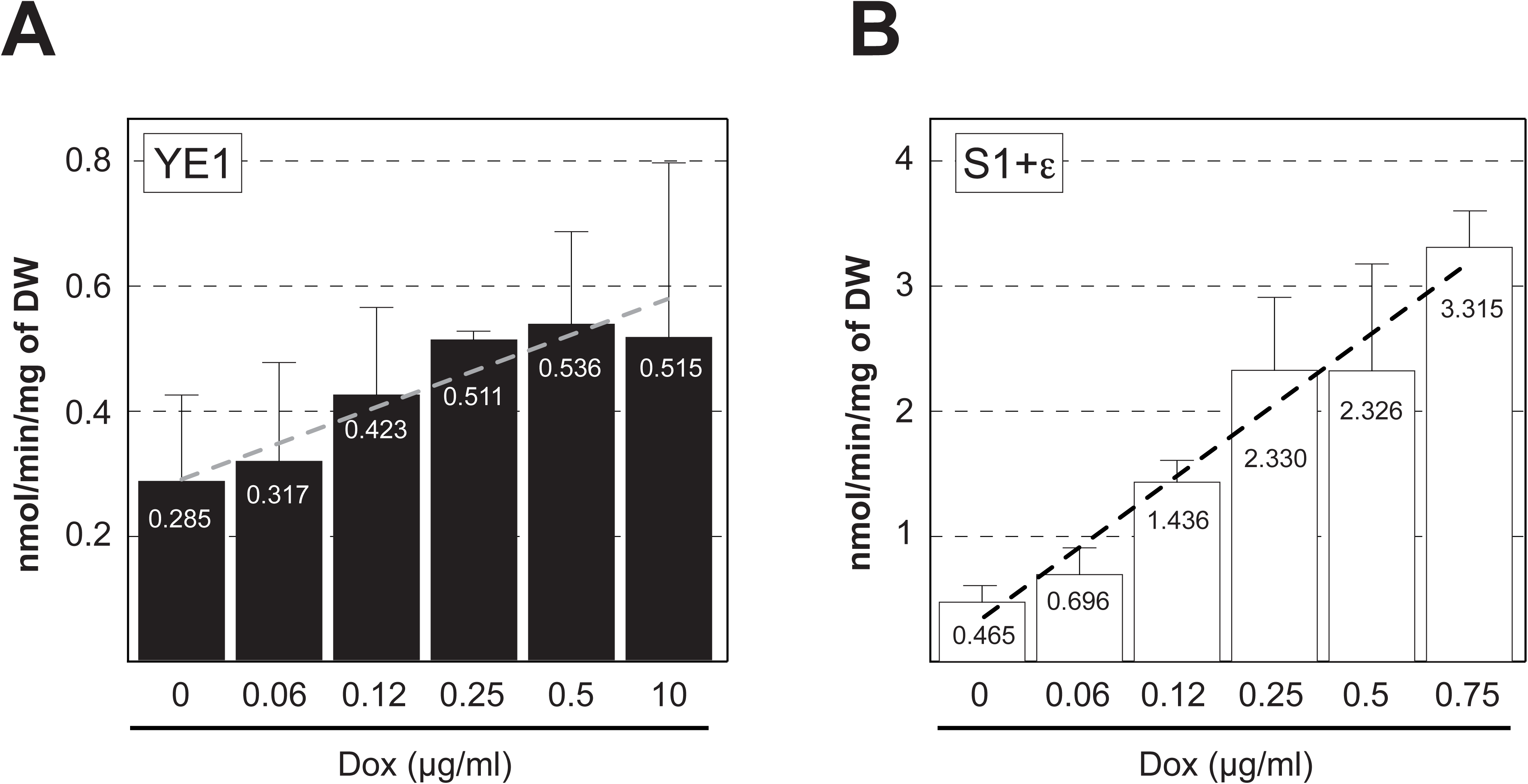
Mitochondrial proliferation. Mitochondrial biogenesis was followed on wild-type YE1 and S1 strains by measuring the activity of the transcription factors HAPs with a β-galactosidase reporter gene, (p161 plasmid, pSDH3-LacZ) in presence of various doxycycline concentrations. DW, dry weight. **(A and B)** YE1 and S1+ε cells respectively were cultivated in glycerol/ethanol condition with various concentration of doxycycline (Dox) that inhibits ε-subunit expression.

**Supplemental Figure 4 (related to Figure 4) -.**
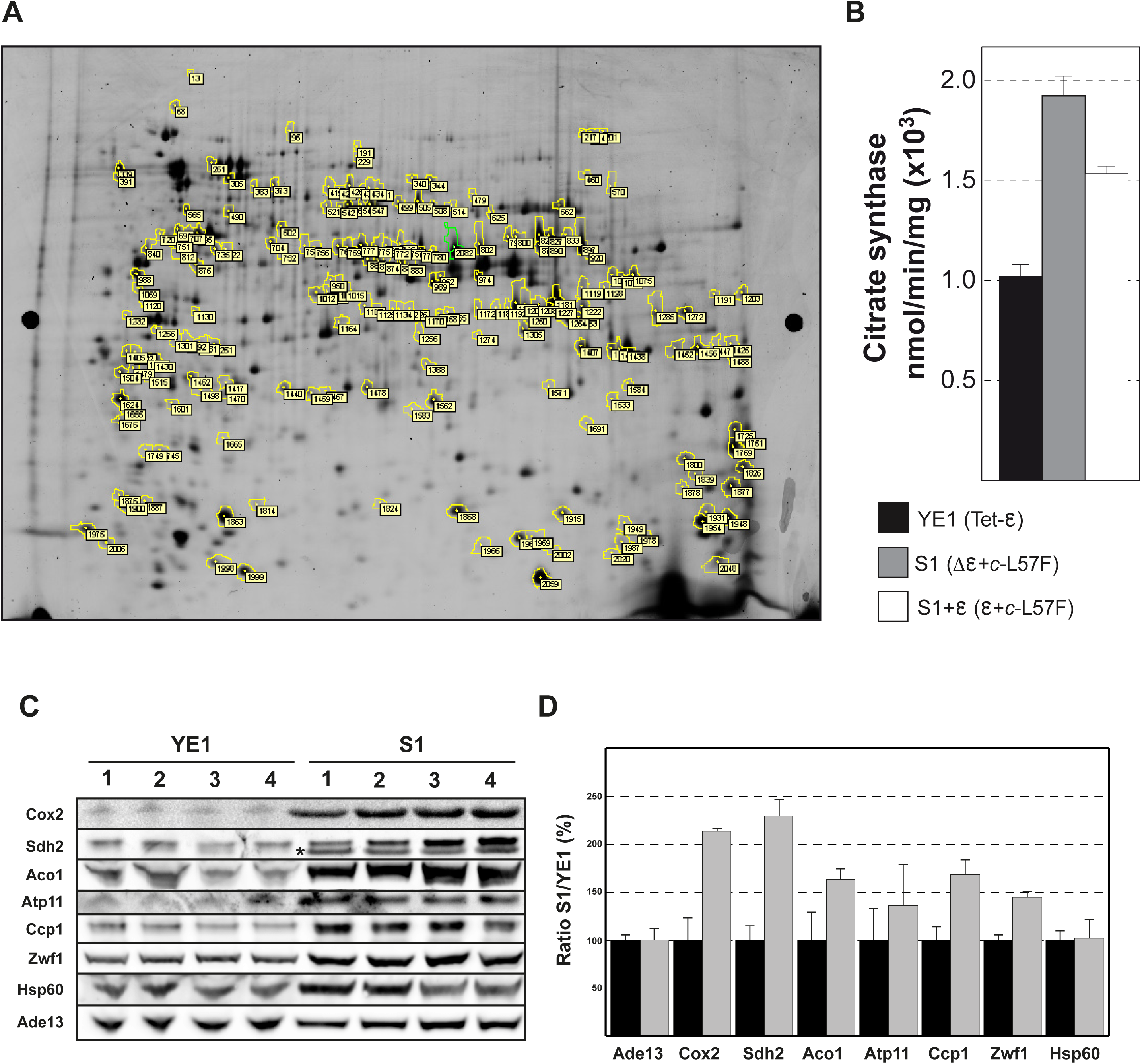
**(A) 2D-DiGE gel.** YE1 protein spot number identified following 2D-DiGE proteomic analysis and listed in Supplemental Table 3. The gel was revealed by silver staining. **(B) Citrate synthase activity.** Citrate synthase activity was performed on isolated mitochondria following the procedure described experimental procedures. **(C/D) Immunoblot and related quantifications of various cytosolic or mitochondrial proteins.** Four different cultures in respiratory condition (glycerol/ethanol) and total cell extracts from wild-type YE1 and S1 mutant strains were analyzed by western-blot. Ade13 was used to normalize the quantifications. Ade13, Adenylosuccinate lyase; Cox2, Subunit 2 of cytochrome-c oxydase; Sdh2, Subunit 2 of succinate dehydrogenase; Aco1, Aconitase; Atp11, ATP synthase chaperone; Ccp1, mitochondrial cytochrome-c peroxidase; Zwf1, Glucose-6-phosphate dehydrogenase; Hsp60, mitochondrial heat-shock protein. The star on the immunoblot indicates the bottom band corresponding to the Sdh2 protein.

**Supplemental Figure 5 (related to Figure 5) -.**
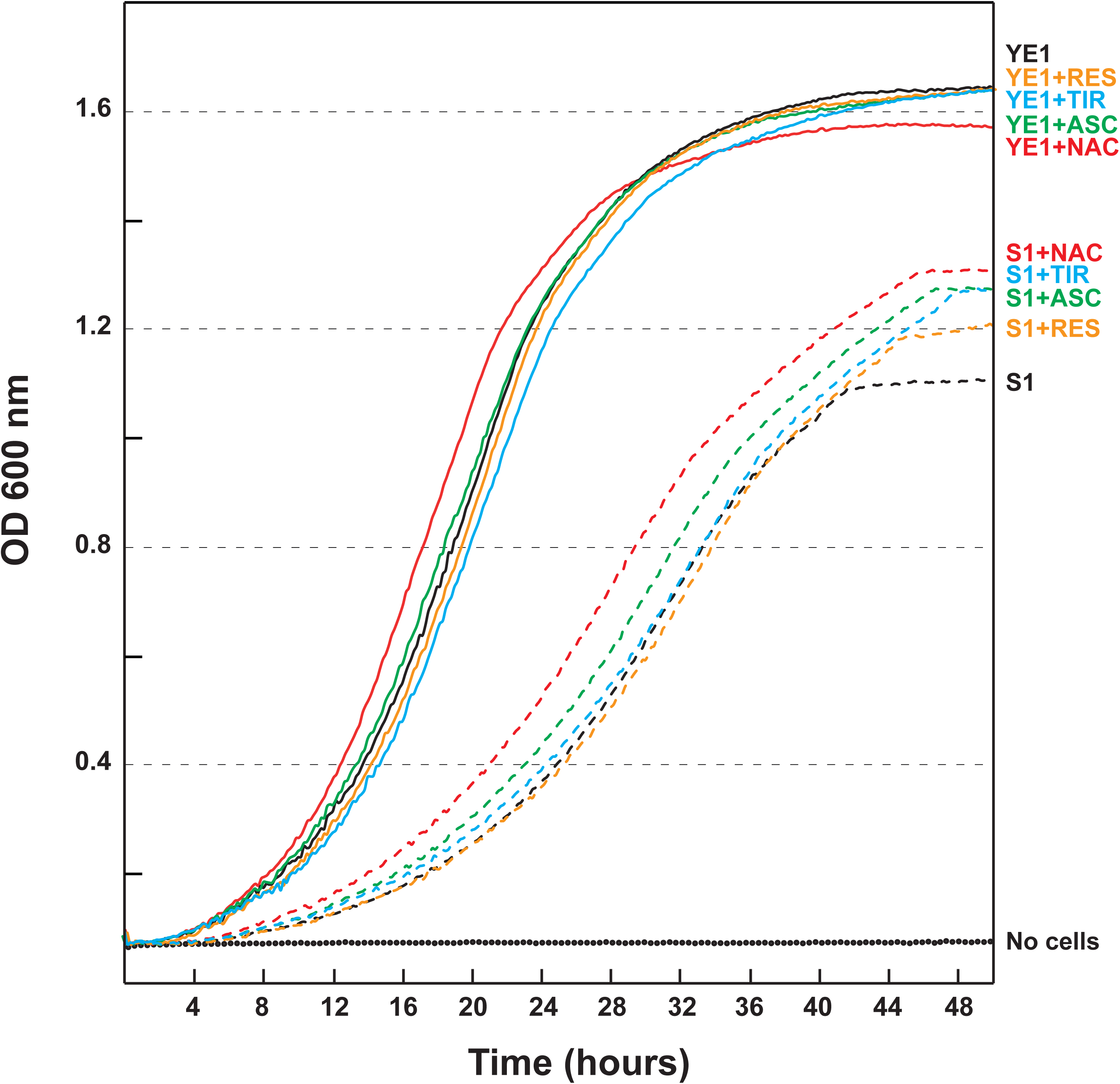
Impact of ROS scavengers on yeast growth. YE1 and S1 cells were grown in glycerol/ethanol condition with various ROS scavengers for 2 days. Optical density (OD 600 nm) was recorded every 10 min at 25°C. N-Acetyl Cystein (NAC, 5 mM, indicated in red); Resveratrol (RES, 100 µM, indicated in orange); Ascorbic acid (ASC, 5 mM, indicated in green) and Tiron (TIR, 5 mM, indicated in blue). The traces are représentative of two experiments.

**Supplemental Figure 6 (related to Figure 6) -.**
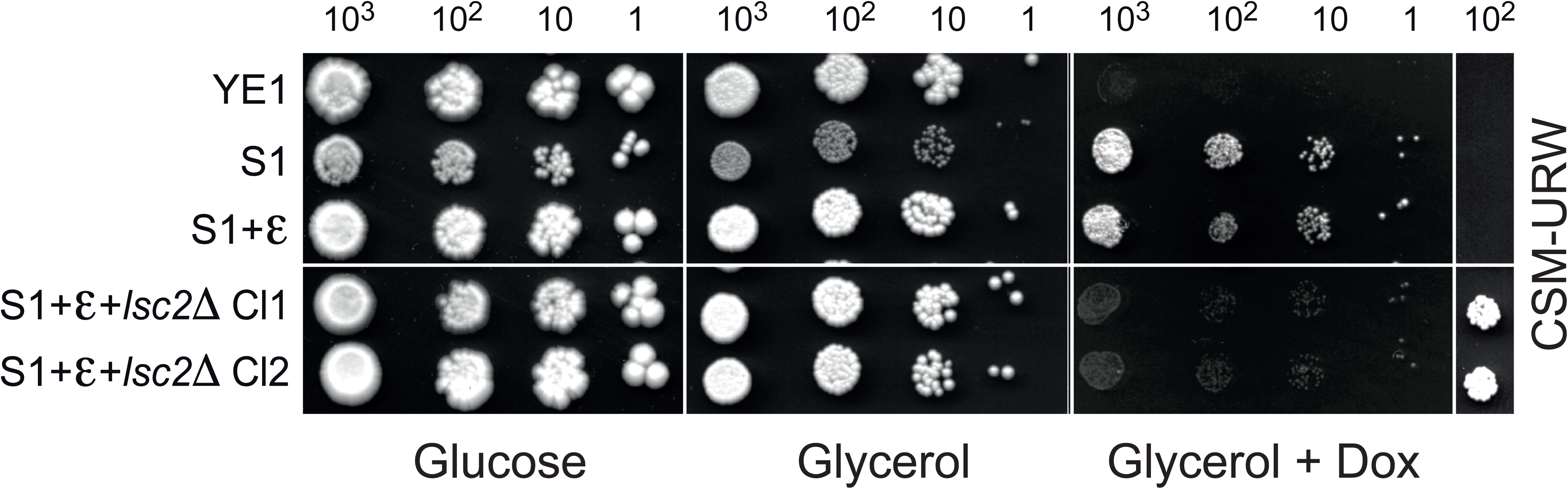
Participation in S1 mutant cells of succinyl-CoA ligase Lsc2 subunit in ATP production. Drop tests of YE1 (Tet-ε), S1 (Δε+c-L57F), S1+ε (ε+c-L57F) and two clones S1+ε+*lsc2*Δ strains on glucose, glycerol, glycerol+doxycycline (10µg/ml) and CSM-URW (complete synthetic medium minus Uracil, Arginine and Tryptophane) media and grown for 3–5 days at 28°C.

**Supplemental Figure 7 (related to Figure 7) -.**
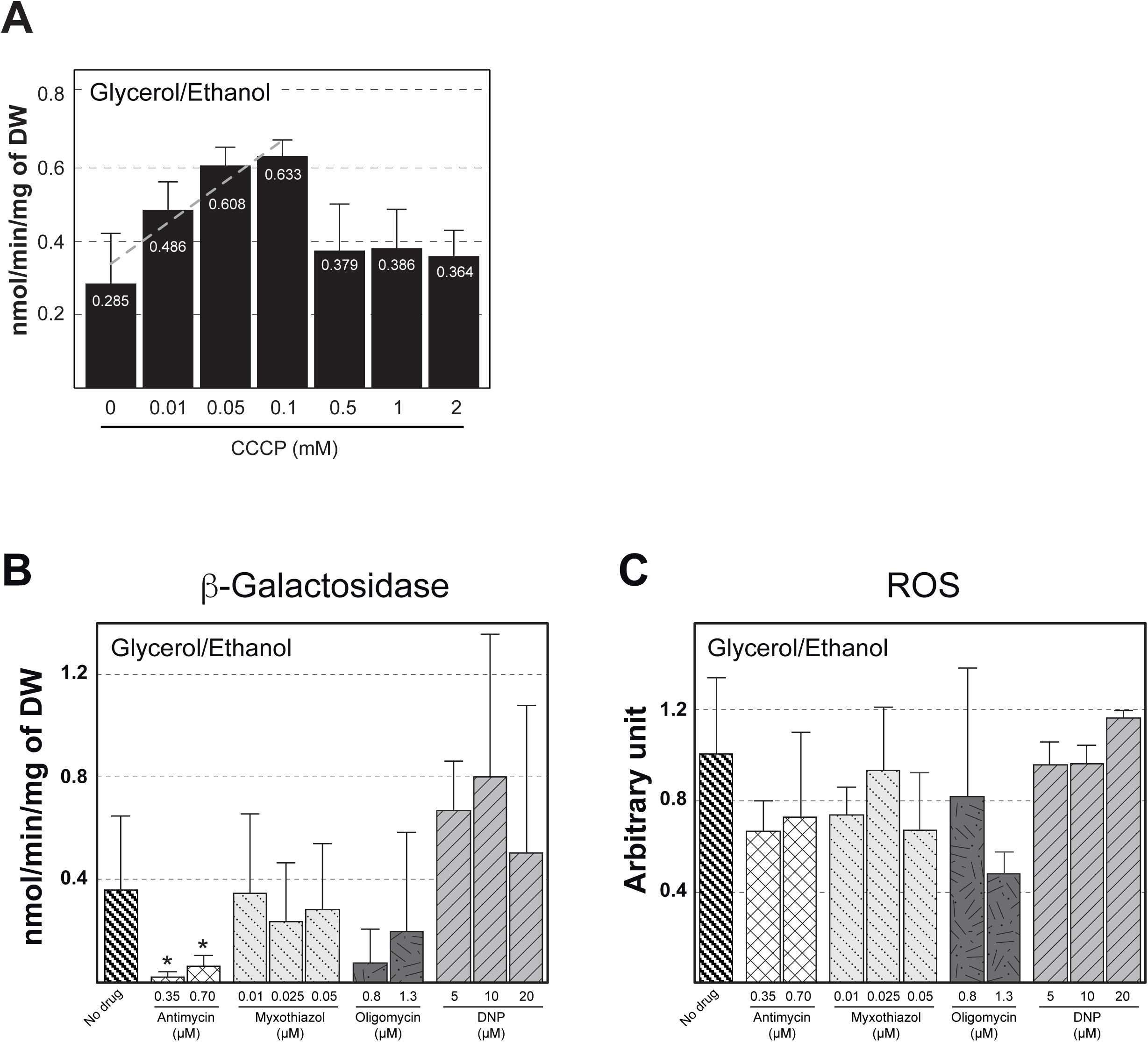
Mitochondrial proliferation and ROS production. **(A)** Mitochondrial biogenesis was followed on wild-type YE1 strain by measuring the activity of the transcription factors HAPs with a β-galactosidase reporter gene, (p161 plasmid, pSDH3-LacZ) in presence of various concentration of the protonophore CCCP. DW, dry weight. n= 5. **(B)** Mitochondrial biogenesis was followed on wild-type W303-1B strain as described previously in presence of various concentration of antimycin (inhibitor of complexe III), myxothiazol (inhibitor of complexe III), oligomycin (inhibitor of complexe V) and 2,4-dinitrophenol (DNP, protonophore). Cells were grown in synthetic 2% glycerol/ethanol medium in absence/presence of inhibitors (4 to 5 generations, OD=2) at 28°C. DW, dry weight. n=3. **(C)** The rate of ROS production in cells was normalized against ROS production of the W303-1B +pSDH3 strain grown in 2% glycerol/ethanol medium without addition of inhibitors. Cells were grown as previoulsy described in B. n=3.

**Table 1 -.** Genotypes of yeast strains.

| Strain name | Nuclear genotype | mtDNA | Source |
| --- | --- | --- | --- |
| W303-1B | *Matα ade2-1 his3-11,15 leu2-3,112 trp1-1 ura3-1 can1-100* | $\rho^+$ | Thomas and Rothstein 1989 |
| W303-1B+pSDH3 | *Matα ade2-1 his3-11,15 leu2-3,112 trp1-1 ura3-1 can1-100 + p161 (LEU2)* | $\rho^+$ | This study |
| W303-1B+lsc1Δ | *Matα ade2-1 his3-11,15 leu2-3,112 trp1-1 ura3-1 can1-100 lsc1Δ::TRP1* | $\rho^+$ | This study |
| YE1 (Tet-ε) | *Matα ade2-1 his3-11,15 leu2-3,112 trp1-1 ura3-1 can1-100 arg8::HIS3 atp15Δ::KanMX + pCM189-ATP15* | $\rho^+ \text{Arg8}^m$ | Tetaud et al. 2014 |
| YE1+pSDH3 | *Matα ade2-1 his3-11,15 leu2-3,112 trp1-1 ura3-1 can1-100 arg8::HIS3 atp15Δ::KanMX + pCM189-ATP15 + p161 (LEU2)* | $\rho^+ \text{Arg8}^m$ | This study |
| YE1+pGFP | *Matα ade2-1 his3-11,15 leu2-3,112 trp1-1 ura3-1 can1-100 arg8::HIS3 atp15Δ::KanMX + pCM189-ATP15 + pFL39-GFP (TRP1)* | $\rho^+ \text{Arg8}^m$ | This study |
| S1 ($\Delta\epsilon$+c-L57F) | *Matα ade2-1 his3-11,15 leu2-3,112 trp1-1 ura3-1 can1-100 arg8::HIS3 atp15Δ::KanMX* | $\rho^+ \text{Arg8}^m$ + c-L57F | Tetaud et al. 2014 |
| S1+pSDH3 | *Matα ade2-1 his3-11,15 leu2-3,112 trp1-1 ura3-1 can1-100 arg8::HIS3 atp15Δ::KanMX + p161 (LEU2)* | $\rho^+ \text{Arg8}^m$ + c-L57F | This study |
| S1+pGFP | *Matα ade2-1 his3-11,15 leu2-3,112 trp1-1 ura3-1 can1-100 arg8::HIS3 atp15Δ::KanMX + pFL39-GFP (TRP1)* | $\rho^+ \text{Arg8}^m$ + c-L57F | This study |
| S1+$\epsilon$ ($\epsilon$+c-L57F) | *Matα ade2-1 his3-11,15 leu2-3,112 trp1-1 ura3-1 can1-100 arg8::HIS3 atp15Δ::KanMX + pCM189-ATP15* | $\rho^+ \text{Arg8}^m$ + c-L57F | Tetaud et al. 2014 |
| S1+$\epsilon$+pSDH3 | *Matα ade2-1 his3-11,15 leu2-3,112 trp1-1 ura3-1 can1-100 arg8::HIS3 atp15Δ::KanMX + pCM189-ATP15 + p161 (LEU2)* | $\rho^+ \text{Arg8}^m$ + c-L57F | This study |
| S1+$\epsilon$+pGFP | *Matα ade2-1 his3-11,15 leu2-3,112 trp1-1 ura3-1 can1-100 arg8::HIS3 atp15Δ::KanMX + pCM189-ATP15 + pFL39-GFP (TRP1)* | $\rho^+ \text{Arg8}^m$ + c-L57F | This study |
| S1+$\epsilon$+lsc1Δ | *Matα ade2-1 his3-11,15 leu2-3,112 trp1-1 ura3-1 can1-100 arg8::HIS3 atp15Δ::KanMX lsc1Δ::TRP1 + pCM189-ATP15* | $\rho^+ \text{Arg8}^m$ + c-L57F | This study |
| S1+$\epsilon$+lsc2Δ | *Matα ade2-1 his3-11,15 leu2-3,112 trp1-1 ura3-1 can1-100 arg8::HIS3 atp15Δ::KanMX lsc1Δ::TRP1 + pCM189-ATP15* | $\rho^+ \text{Arg8}^m$ + c-L57F | This study |

**Table 2 –.**
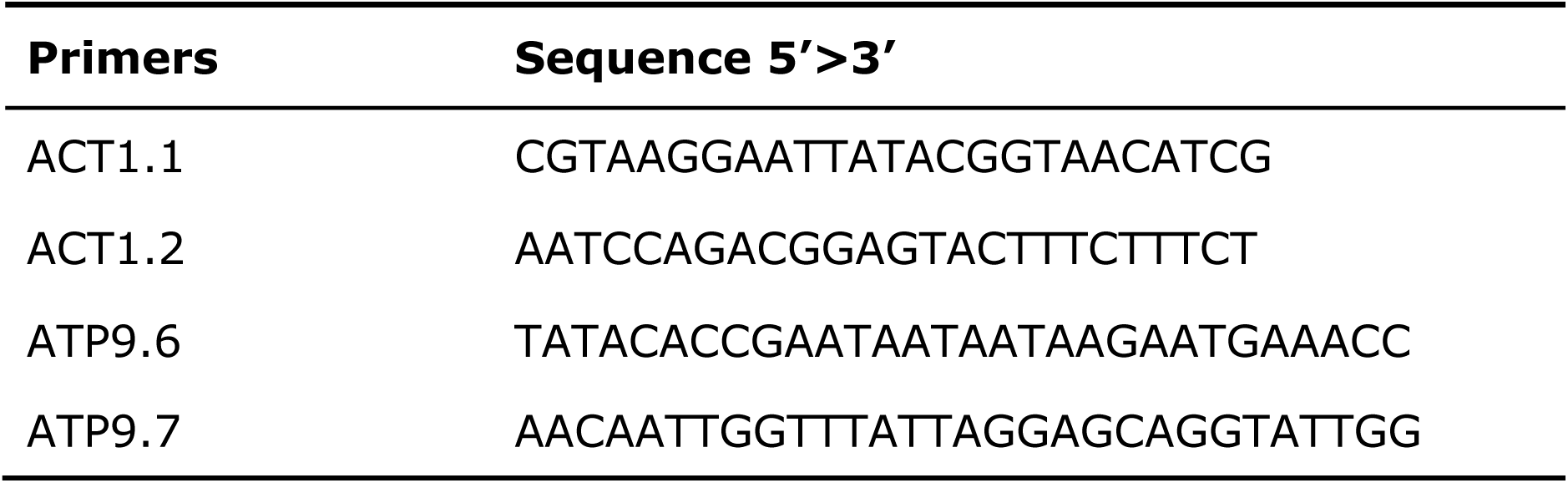
qPCR primers.

**Supplemental Table 3:**
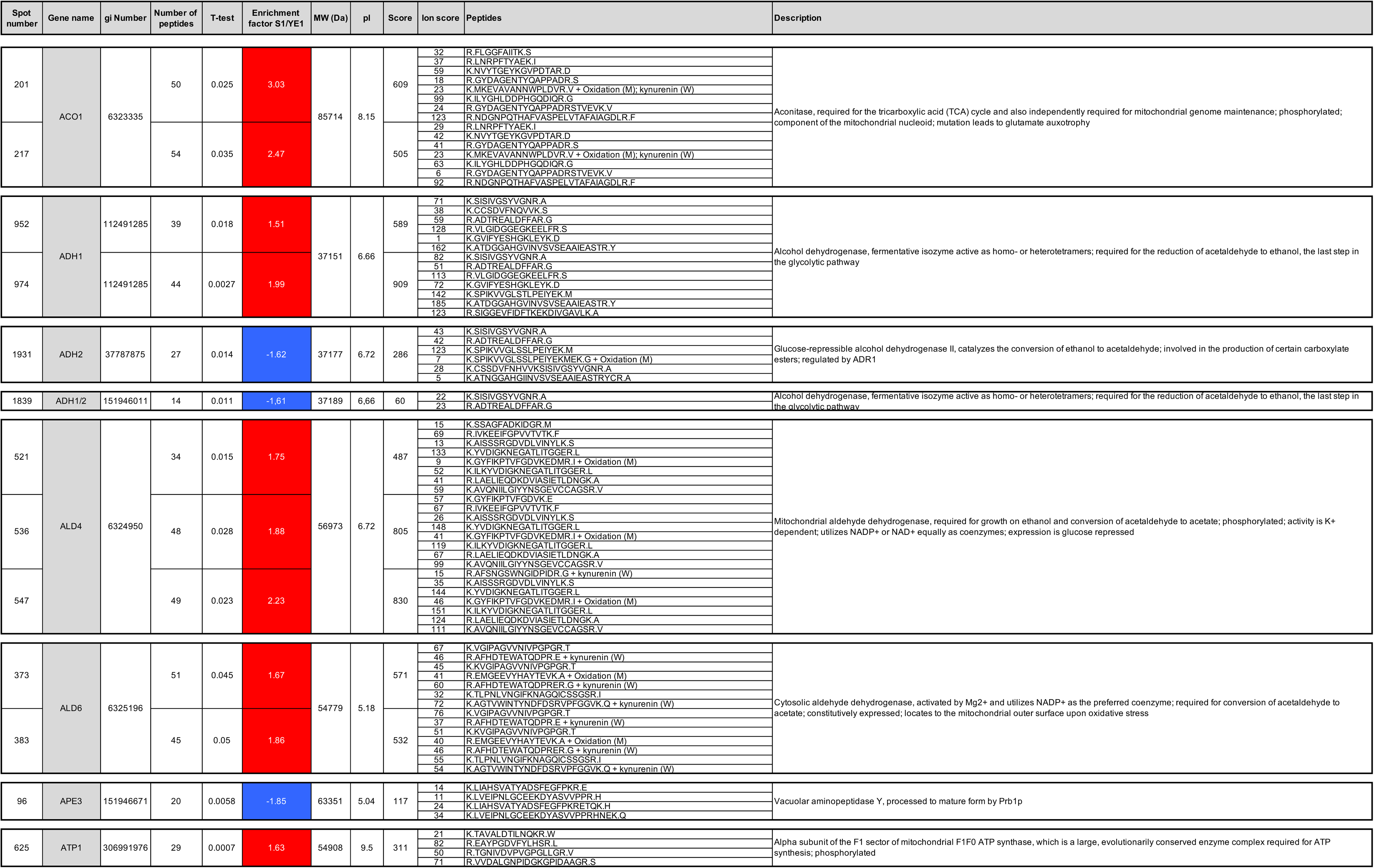
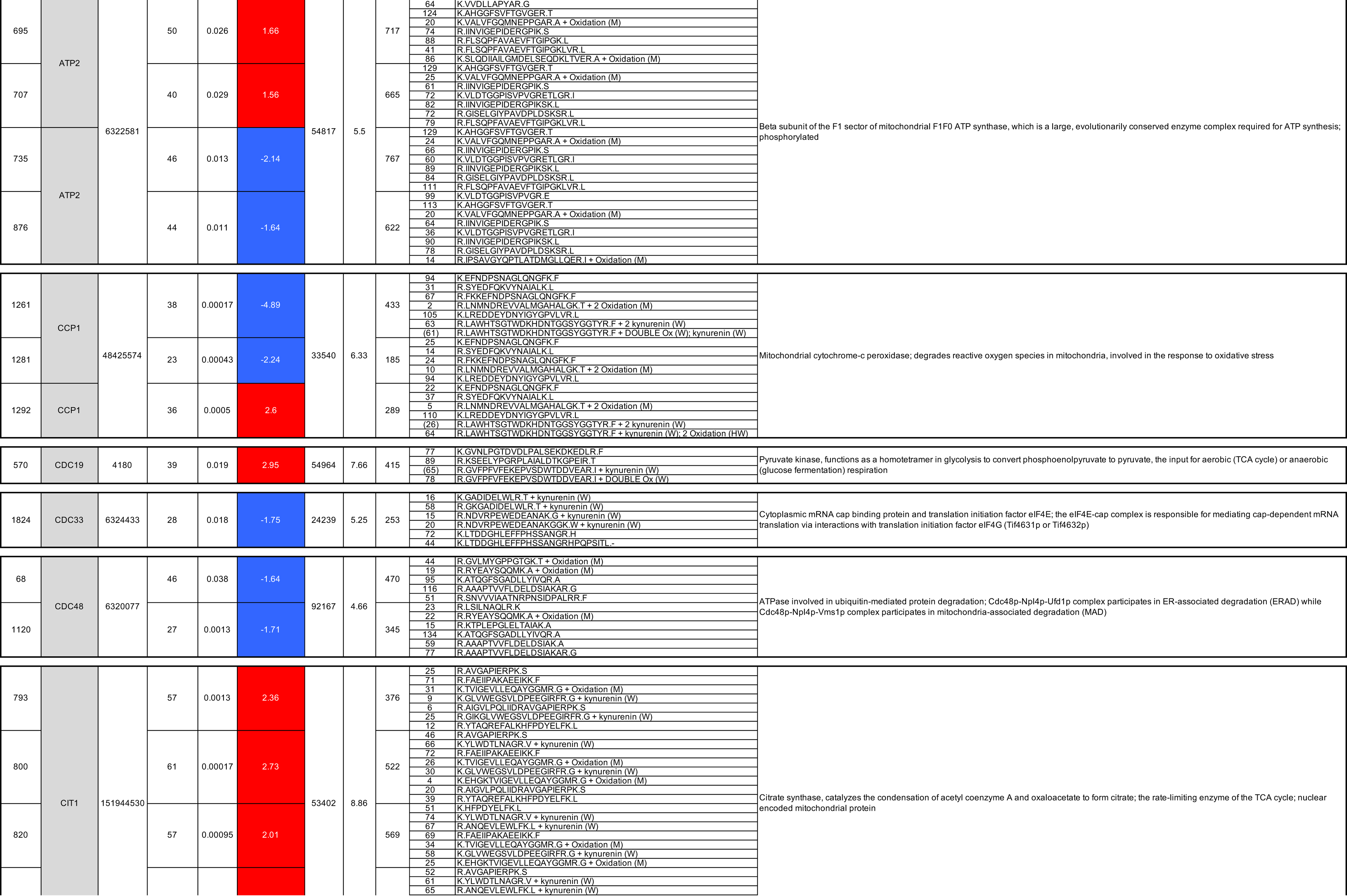
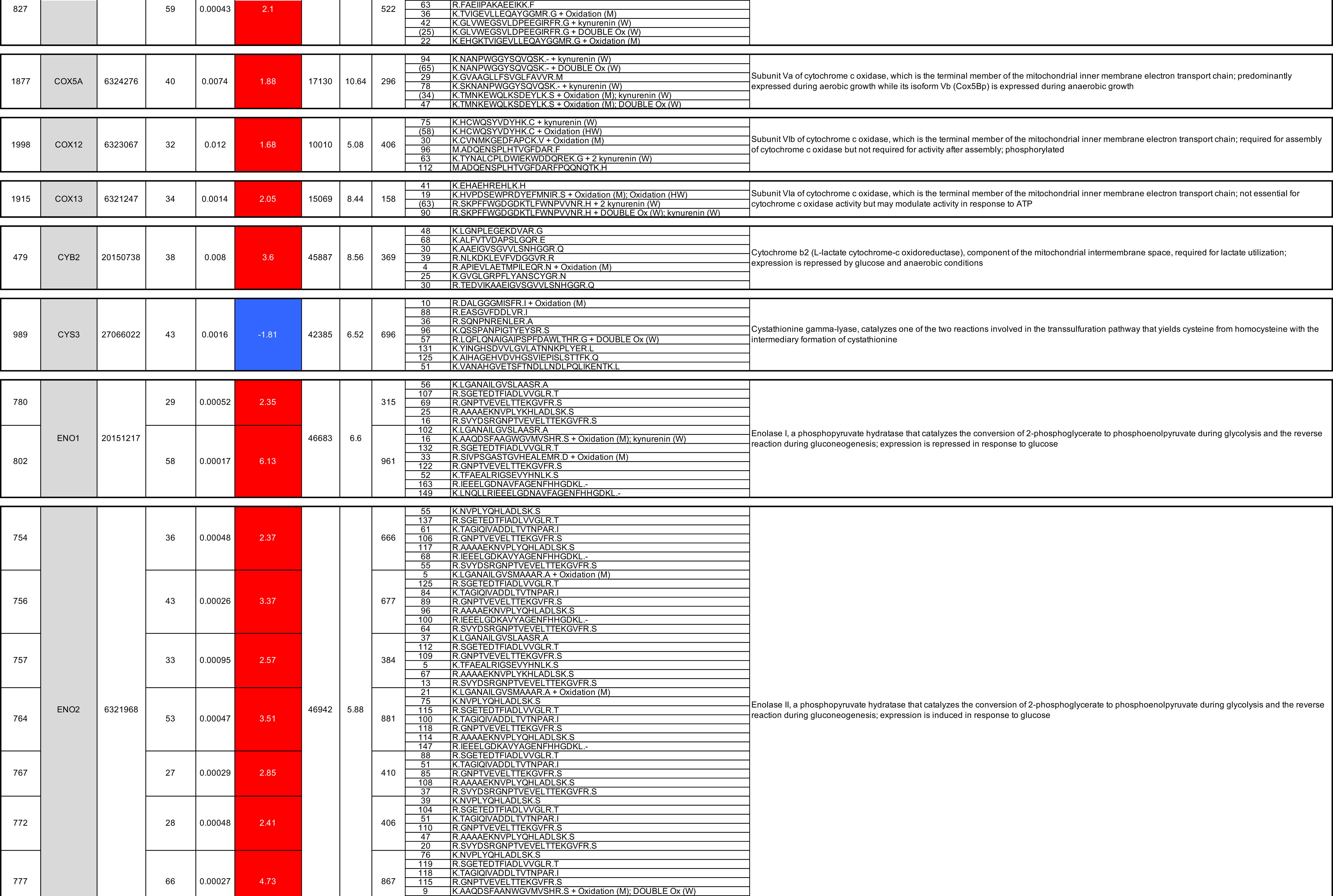
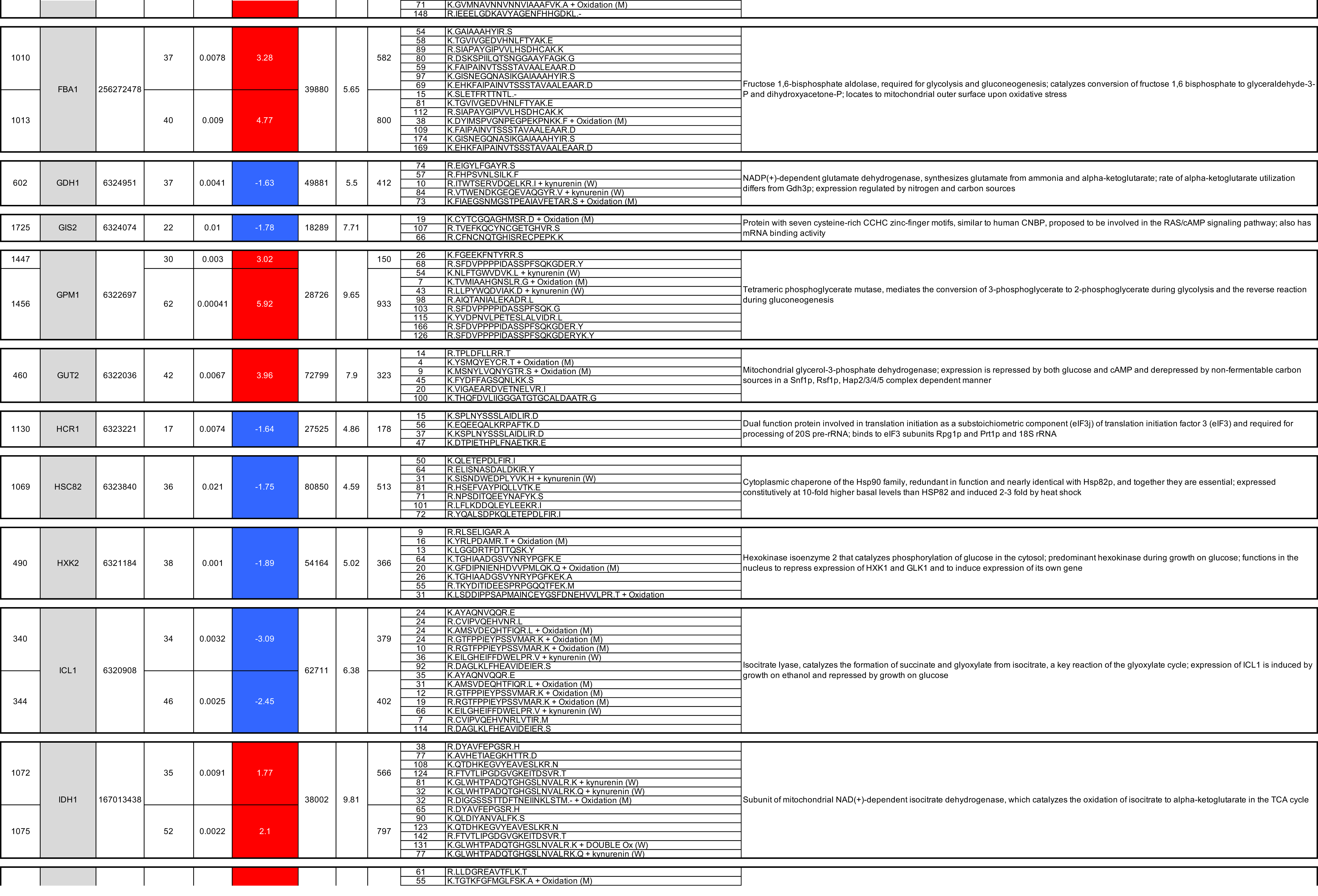
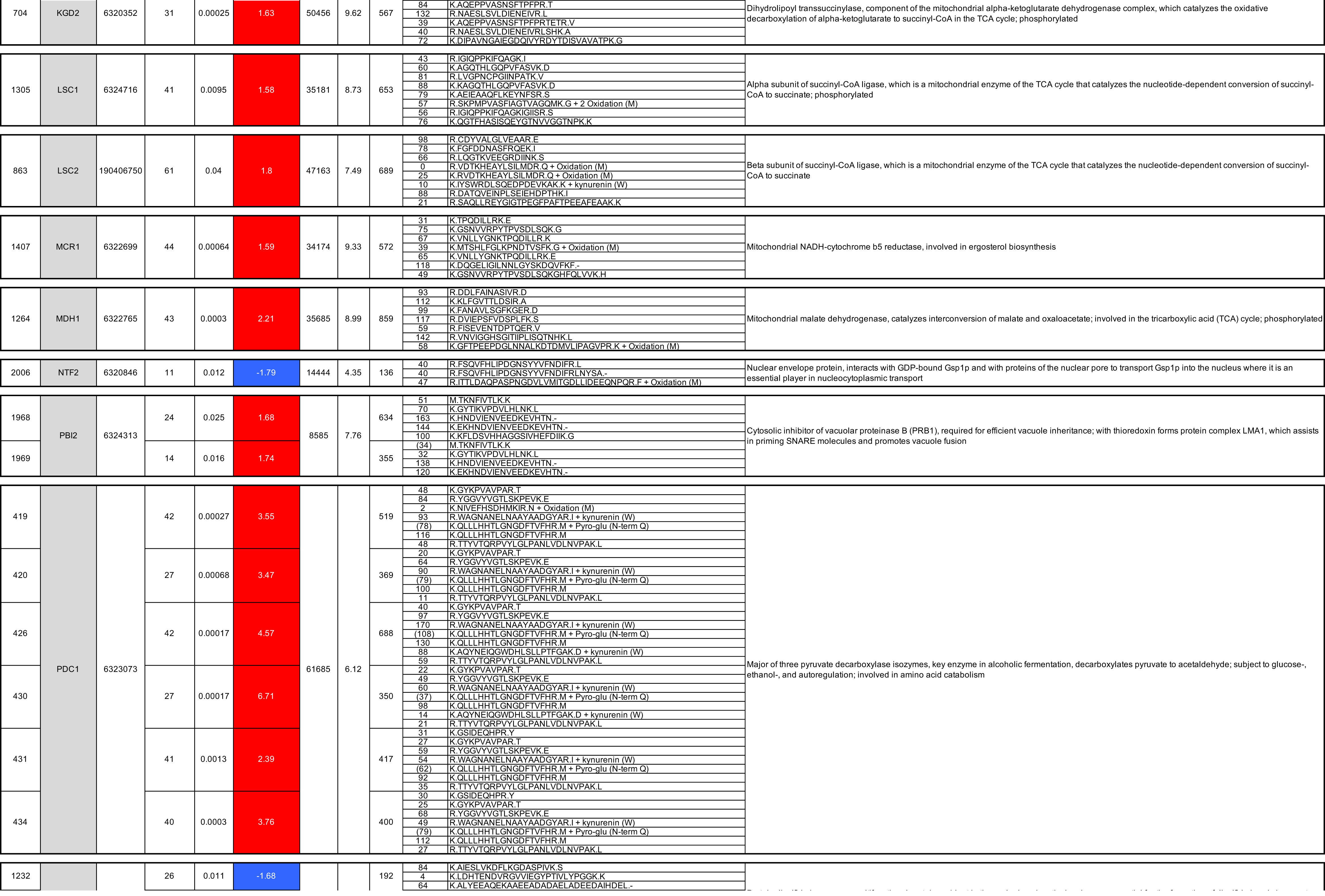
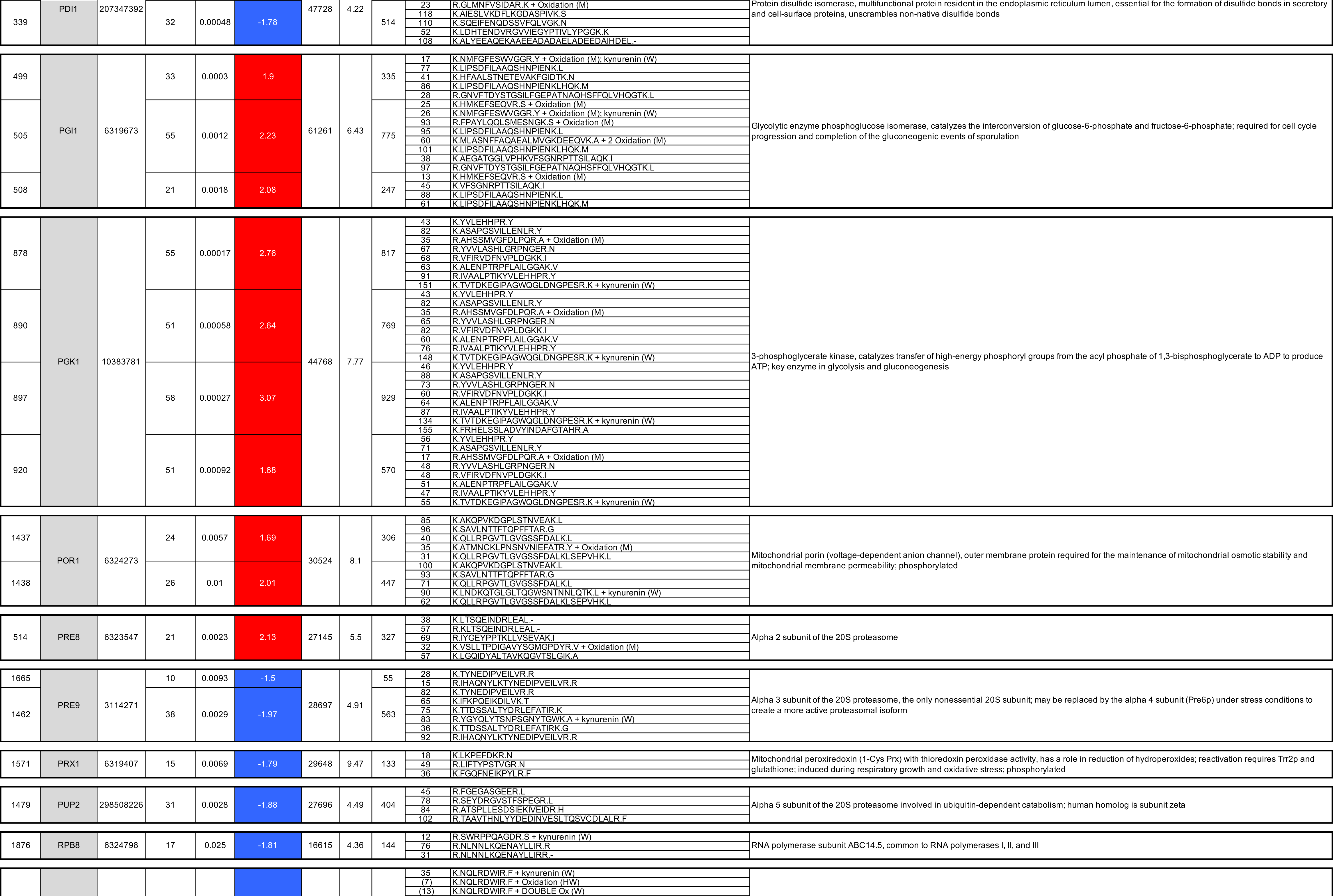
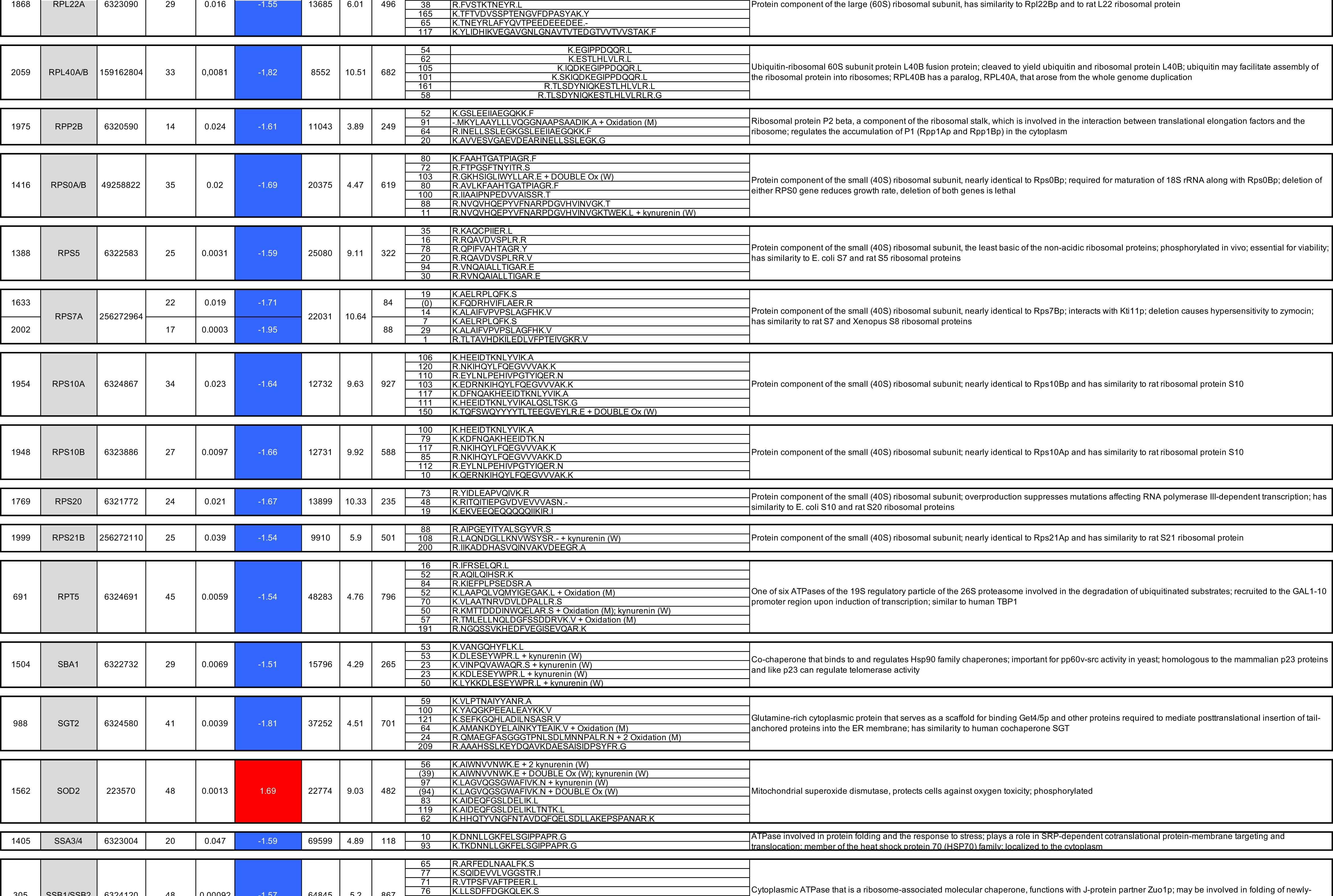
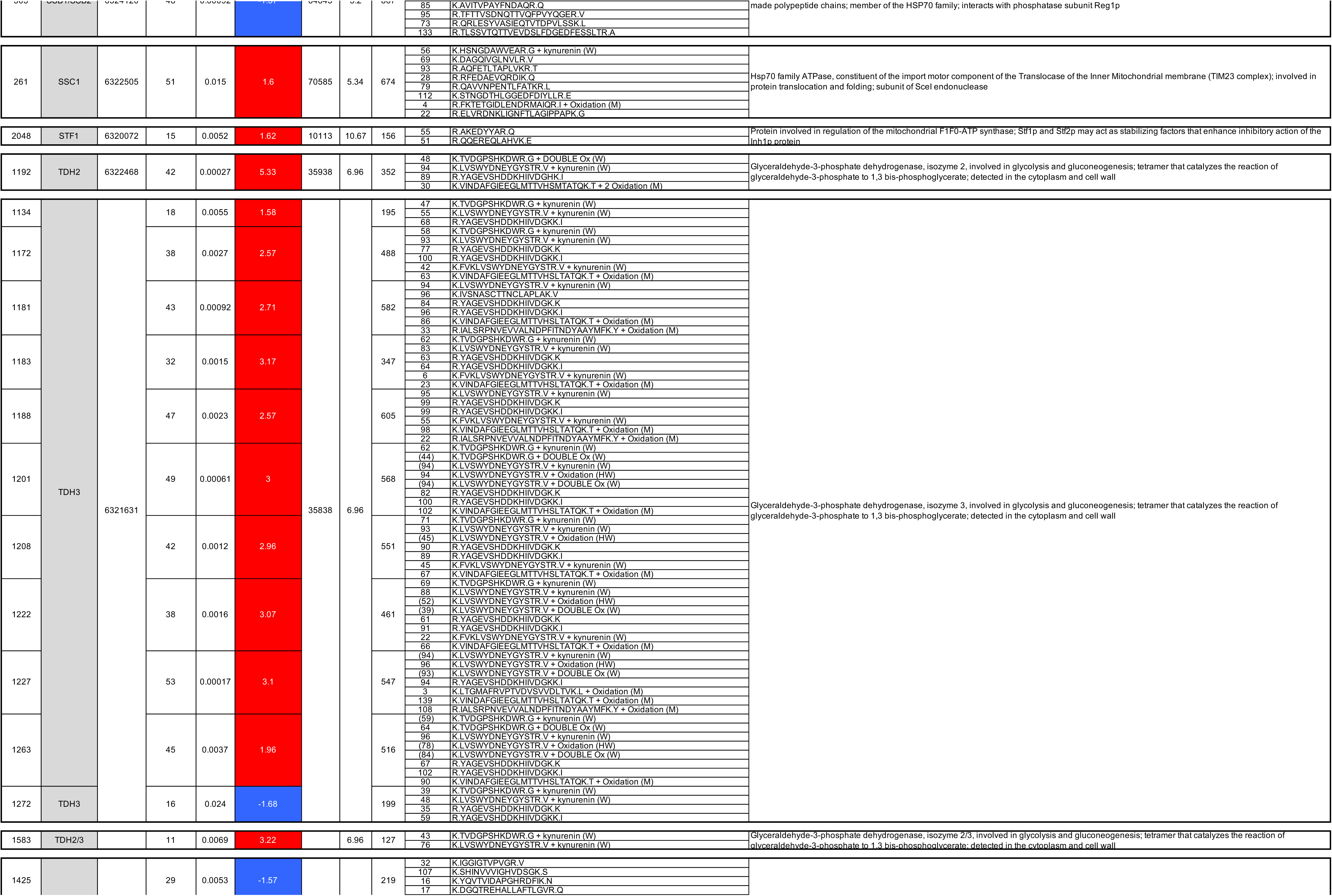
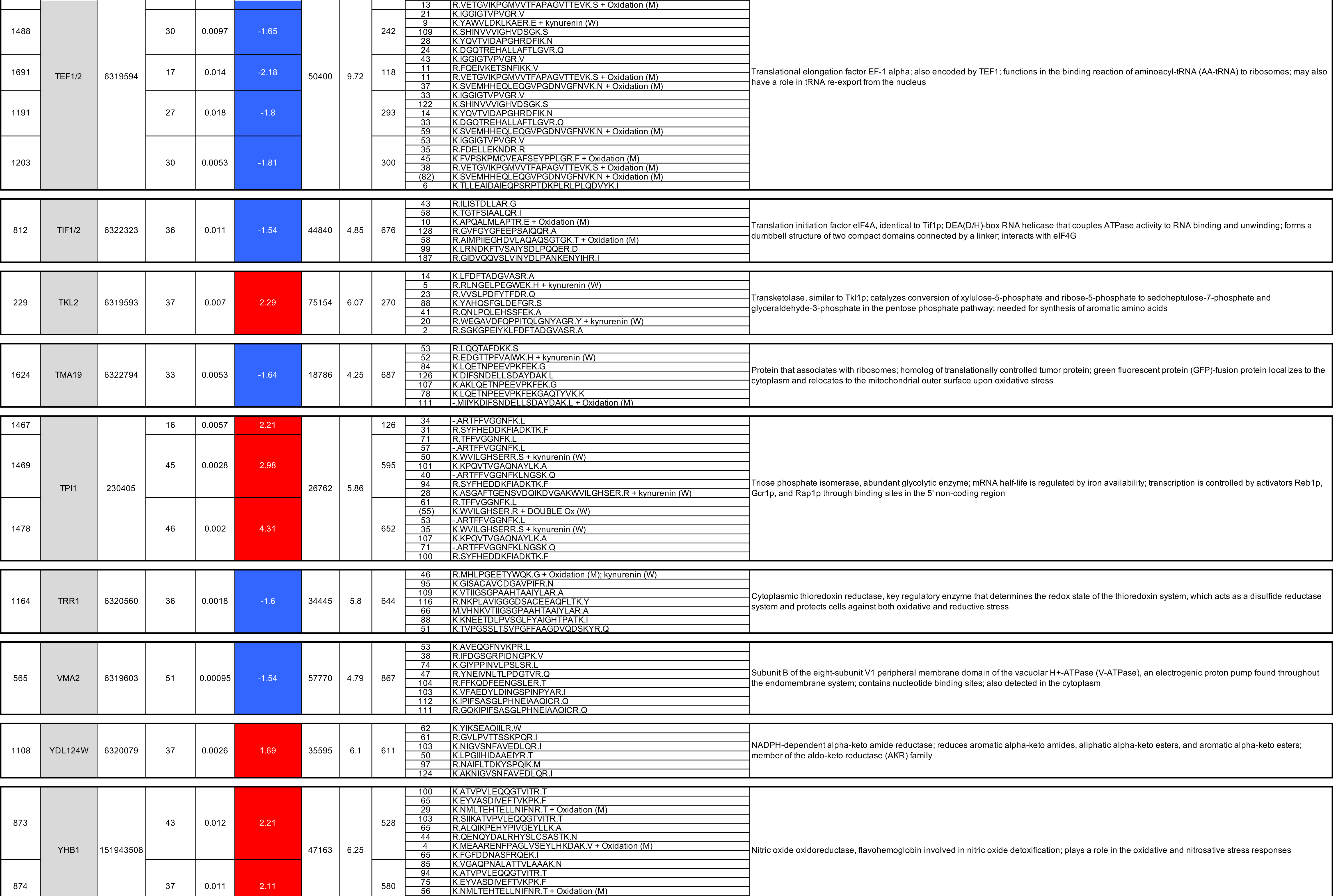
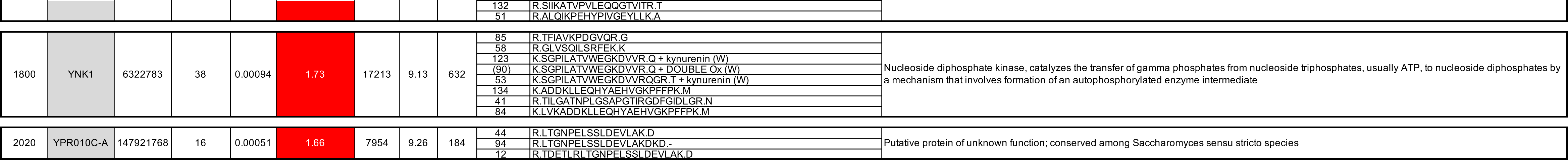
Spot number, protein spot number identified following 2D-DiGE presented in Supplemental Figure 3. gi Number, identified according to GenBank. Number of peptides, number of peptide masses matched the top hit from MS-Fit PMF (peptide mass fingerprint). Enrichment factor S1/YE1, positive value means that the protein was up regulated in S1 strain compared to YE1 strain and was indicated in red; negative value means that the protein was down regulated in S1 strain compared to YE1 strain and was indicated in blue. MW, predicted molecular weight in Dalton according to protein sequence. pI, predicted isoelectric point of the protein according to sequence. Score, Mascot MS protein score obtained by MALDI-ToF/ToF spectra. Ion score, Mascot MS/MS ion score obtained by MALDI-ToF/ToF spectra.

